# Clustered γ-Protocadherins Regulate Cortical Interneuron Programmed Cell Death

**DOI:** 10.1101/2020.01.14.906941

**Authors:** Walter Mancia, Julien Spatazza, Benjamin Rakela, Ankita Chatterjee, Viraj Pande, Tom Maniatis, Andrea R. Hasenstaub, Michael P. Stryker, Arturo Alvarez-Buylla

**Author notes:** These authors contributed equally (Walter Mancia, Julien Spatazza).

## Abstract

Cortical function critically depends on inhibitory/excitatory balance. GABAergic cortical inhibitory interneurons (cINs) are born in the ventral forebrain. After completing their migration into cortex, their final numbers are adjusted-during a period of postnatal development - by programmed cell death (PCD). The mechanisms that regulate cIN elimination remain controversial. Here we show that genes in the protocadherin (Pcdh)-γ gene cluster, but not in the Pcdh-α or Pcdh-β clusters, are required for survival of cINs through a BAX-dependent mechanism. Surprisingly, the physiological and morphological properties of Pcdh-γ deficient and wild type cINs during PCD were indistinguishable. Co-transplantation of wild type and Pcdh-γ deficient interneuron precursor cells demonstrate that: 1) the number of mutant cINs eliminated was much higher than that of wild type cells, but the proportion of mutant or WT cells undergoing cell death was not affected by their density; 2) the presence of mutant cINs increases cell death among wild-type counterparts, and 3) cIN survival is dependent on the expression of Pcdh-γ C3, C4, and C5. We conclude that Pcdh-γ, and specifically γC3, γC4, and γC5, play a critical role in regulating cIN survival during the endogenous period of PCD.

**Significance:** GABAergic cortical inhibitory interneurons (cINs) in the cerebral cortex originate from the ventral embryonic forebrain. After a long migration, they come together with local excitatory neurons to form cortical circuits. These circuits are responsible for higher brain functions, and the improper balance of excitation/inhibition in the cortex can result in mental diseases. Therefore, an understanding of how the final number of cINs is determined is both biologically and, likely, therapeutically significant. Here we show that cell surface homophilic binding proteins belonging to the clustered protocadherin gene family, specifically three isoforms in the Pcdh-γ cluster, play a key role in the regulation cIN programmed cell death. Co-transplantation of mutant and wild-type cINs shows that Pcdh-γ genes have cell-autonomous and non-cell autonomous roles in the regulation of cIN cell death. This work will help identify the molecular mechanisms and cell-cell interactions that determine how the proper ratio of excitatory to inhibitory neurons is determined in the cerebral cortex.

## Introduction

GABAergic cortical inhibitory interneurons (cINs) regulate neuronal circuits in the neocortex. The ratio of inhibitory interneurons to excitatory neurons is crucial for establishing and maintaining proper brain functions (J. L. R. Rubenstein, 2003; Huang, Di Cristo and Ango, 2007; Chao *et al*., 2010; Rossignol, 2011; Marín, 2012; Hattori *et al*., 2017). Alterations in the number of cINs have been linked to epilepsy (F. Edward Dudek, 2003), schizophrenia (Beasley and Reynolds, 1997; Hashimoto *et al*., 2003; Enwright *et al*., 2016) and autism (Fatemi *et al*., 2009; Giada Cellot, 2014; Gao and Penzes, 2015). During mouse embryonic development, the brain produces an excess number of cINs, and ∼ 40% of those are subsequently eliminated by apoptosis during early postnatal life, between postnatal day (P)1 and 15 (Southwell *et al*., 2012; Denaxa, Neves, Burrone, *et al*., 2018; Wong *et al*., 2018). What makes the death of these cells intriguing is its timing and location. In normal development, cINs are generated in the medial and caudal ganglionic eminences (MGE; CGE) of the ventral forebrain, far from their final target destination in the cortex. cINs migrate tangentially from their sites of birth to reach the neocortex where they become synaptically integrated and complete their maturation (Anderson *et al*., 1997; Wichterle *et al*., 2001; Nery, Corbin and Fishell, 2003; Butt *et al*., 2005). The ganglionic eminences are also an important source of interneurons in the developing human brain, where migration and differentiation extend into postnatal life (Hansen *et al*., 2013; Ma *et al*., 2013; Paredes *et al*., 2016). How is the final number of cINs is regulated once these cells arrive in the cortex?

Since cINs play a pivotal role in regulating the level of cortical inhibition, the adjustment of their number by programmed cell death is a key feature of their development and essential for proper brain physiology. While recent work suggests that activity-dependent mechanisms regulate cIN survival through their connectivity to excitatory neurons (Denaxa, Neves, Burrone, *et al*., 2018; Wong *et al*., 2018; Duan *et al*., 2020), (Denaxa, Neves, Rabinowitz, *et al*., 2018) studies indicate that cIN survival is mediated by a population - or self-autonomous mechanism (Rauskolb *et al*., 2010; Southwell *et al*., 2012). Indeed, heterochronically transplanted MGE cIN precursors display a wave of apoptosis coinciding with their age, and asynchronously from endogenous cINs. Whereas it is well established that neuronal survival in the peripheral nervous system (PNS) is regulated through limited access to neurotrophic factors secreted by target cells (Eric J Huang, 2001; Luigi Aloe, 2013; Oppenheim, Milligan and von Bartheld, 2013)), cIN survival is independent of TrkB, the main neurotrophin receptor expressed by neurons of the CNS (Rauskolb *et al*., 2010; Southwell *et al*., 2012).

Moreover, the proportion of cINs undergoing apoptosis remains constant across graft sizes that vary 200-fold (Rauskolb *et al*., 2010; Southwell *et al*., 2012). Altogether, this work suggests that cIN developmental death is intrinsically determined and that cell-autonomous mechanisms within the maturing cIN population contribute to the regulation of their own survival.

The clustered protocadherins (Pcdh) (Wu and Maniatis, 1999) are a set of cell surface homophilic binding proteins implicated in neuronal survival and self-avoidance in the spinal cord, retina, cerebellum, hippocampus and glomerulus (Wang *et al*., 2002; Lefebvre *et al*., 2008, 2012; Chen *et al*., 2017; Katori *et al*., 2017; Mountoufaris *et al*., 2017; Ing-Esteves *et al*., 2018). In the mouse, the Pcdh locus encodes a total of 58 isoforms that are arranged in three gene clusters: Pcdh-α, Pcdh-β, and Pcdh-γ ((Wu *et al*., 2001). The Pcdh-α and Pcdh-γ isoforms are each composed of a set of variable exons, which are spliced to three common constant cluster-specific exons (Tasic *et al*., 2002; Wang, 2002). Each variable exon codes for the extracellular, transmembrane and most-proximal intracellular domain of a protocadherin protein. The Pcdh-β isoforms are encoded by single exon genes encoding both extracellular, transmembrane and cytoplasmic domains (Wu and Maniatis, 1999). Of the 58 Pcdh genes, it has been suggested that a combinatorial, yet stochastic, set of isoforms is expressed in each neuron (Esumi *et al*., 2005; Kaneko *et al*., 2006; Mountoufaris *et al*., 2017), suggesting a source for neuronal diversity in the CNS (Canzio *et al*., 2019).

Interestingly, Pcdh-γ genes, and specifically isoforms γC3, γC4, and γC5, are required for postnatal survival in mice (Wang *et al*., 2002; Chen *et al*., 2012; Hasegawa *et al*., 2016). Whether Pcdh genes are required for the regulation of cIN elimination remains unknown.

In the present study, we used a series of genetic deletions of the Pcdh gene locus to probe the role of clustered Pcdhs in the regulation of cIN cell death in mice. We show that Pcdh-γ, but not Pcdh-α or Pcdh-β, are required for the survival of approximately 50% of cINs through a BAX-dependent mechanism. Using co-transplantation of Pcdh-γ deficient and wild-type (WT) cells of the same age, we show that cINs compete for survival in a mechanism that involves Pcdh-γ. Taking advantage of the transplantation assay, we show that removal of the three Pcdh-γ isoforms, γC3, γC4, and γC5, is sufficient to increase cell death of MGE-derived cINs. Three-dimensional reconstructions and patch-clamp recordings indicate that the Pcdh-γ mutant cells have similar morphology, excitability and receive similar numbers of inhibitory and excitatory synaptic inputs compared to wild type cINs. We conclude that cIN cell death is regulated by all or some of the C isoforms in the Pcdh-γ cluster and that this process is independent of the structural complexity or physiological state of the cell.

## Results

### Pcdh-γ expression in developing cINs

Expression of clustered protocadherins (Pcdh) in the brain starts in the embryo and continues postnatally (Kohmura *et al*., 1998; Wang *et al*., 2002; Frank *et al*., 2005; Hirano *et al*., 2012). RT-PCR analysis revealed the expression of each of the 58 isoforms in the Pcdh gene locus in the adult cortex (P30) (**Figure 1A**). Of the 58 Pcdh genes, those in the Pcdh-γ cluster are essential for postnatal survival (Chen *et al*., 2012; Hasegawa *et al*., 2016), and are implicated in cell death in the retina and spinal cord (Lefebvre *et al*., 2008; Prasad *et al*., 2008). We, therefore, determined whether Pcdh-γ genes are expressed in cINs during the period of cIN cell death. Using *Gad67-GFP* mice to label GABAergic cINs (Tamamaki *et al*., 2003), we FACS-sorted GFP-positive (GFP+) and GFP-negative (GFP-) cells from P7 mice, at the peak of cIN cell death (**Figure 1- Figure supplement 1A**). We confirmed that GABAergic cell markers (Gad1, Gad2) were enriched in the GFP+ population, while markers of excitatory neurons (Tbr1, Satb2, Otx1), astrocytes (GFAP, Aldh1L1), and oligodendrocytes (Olig2, MBP) were enriched in the GFP-population (**Figure 1- Figure supplement 1B**).

**Figure 1.**
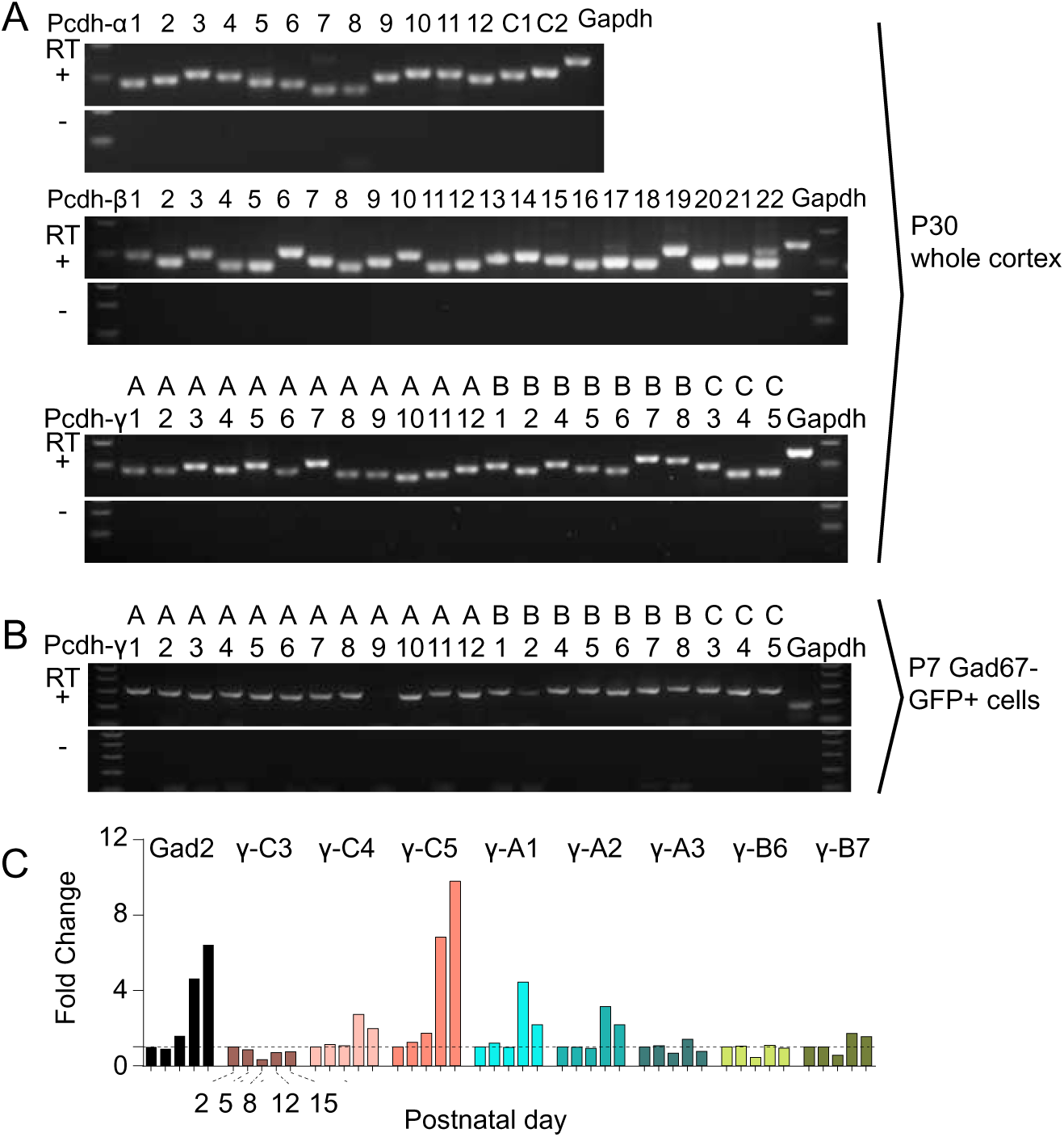
Expression of clustered Pcdhs in the mouse cortex and purified cortical GAB-Aergic cells. **A.** PCR analysis of clustered Pcdh and GAPDH gene expression in P30 whole cortex extracts. **B.** PCR analysis of Pcdh-γ and GAPDH gene expression in purified P7 cortical GABAergic cells. **C.** Quantification of target gene mRNA levels at various postnatal stages (P2, P5, P8, P12, P15) in purified cortical GABAergic cells. P2 mRNA levels used as a reference for each gene

With the exception of A9 Pcdh-γ isoform, we detected expression of all other 21 Pcdh-γ (RT-PCR) in cINs (**Figure 1B**). To determine the expression pattern of Pcdh-γ at different stages during the period of cell death, we measured the expression level of 8 Pcdh-γ mRNAs (γC3-5, γA1-A3, γB6-7) at P2, P5, P8, P12 and P15 using qPCR (**Figure 1C**). All 8 isoforms were expressed in cINs at each of the 5 ages studied. Interestingly, the expression of γ-C5 increased dramatically between P8 and P15. An increase in expression of γA1, γA2, and γC4 was also observed at P12, compared to other ages, but this increase was less pronounced than that observed for γC5. The above results show that all Pcdh isoforms are expressed in cINs and that the expression of γA1, γA2, γC4, and γC5 increases during the period of postnatal cell death.

### Reduced number of cIN in the cortex of Pcdh-γ mutants

Most cINs are produced between E10.5 and E16.5 by progenitors located in the medial and caudal ganglionic eminences (MGE and CGE) (Anderson *et al*., 1997; Wichterle *et al*., 2001; Nery, Fishell and Corbin, 2002; Miyoshi *et al*., 2010). To address the potential role of Pcdh-γ in cIN development, we used the Pcdh-γ conditional allele (*Pcdh-γ^fcon3^*) to block production of all 22 Pcdh-γ isoforms (Lefebvre *et al*., 2008). In the *Pcdh-γ^fcon3^* allele, the third common exon shared by all *Pcdh-γ* isoforms contains the sequence coding for GFP and is flanked by loxP sites (Lefebvre *et al*., 2008)(**Figure 2A**). In unrecombined Pcdh-γ^fcon3^ mice, all Pcdh-γ isoforms are thus fused to GFP. However, when these animals are crossed to a Cre driver line, expression of the entire Pcdh-γ cluster is abolished in Cre-expressing cells (Prasad *et al*., 2008). Robust GFP expression was detected throughout the brain in E13.5 embryos, including cells in the MGE and CGE (**Figure 3B**), indicating expression of Pcdh-γ isoforms in cIN progenitors. We crossed *Pcdh-γ^fcon3^* mice to Gad2^Cre^ mice (Taniguchi *et al*., 2011) to conditionally ablate all Pcdh-γ in GABAergic cells throughout the CNS at an early embryonic stage (E10.5)(Katarova *et al*., 2000). Recombined cells were visualized thanks to the conditional tdTomato reporter allele Ai14 (**Figure 2A**). Heterozygous *Gad2^cre^; Ai14; Pcdh-γ^fcon3/^*^+^ mice were viable and fertile. However, homozygous *Gad2^cre^; Ai14; Pcdh-γ^fcon3/fcon3^* mice displayed growth retardation after birth, a hind limb paw-clasping phenotype when held by the tail and were infertile (**Figure 2B**). Brain size as well as cerebral cortex thickness of homozygous *Gad2^cre^; Ai14; Pcdh-γ^fcon3/fcon3^* was similar to those of control mice (**Figure 2B’**).

**Figure 2.**
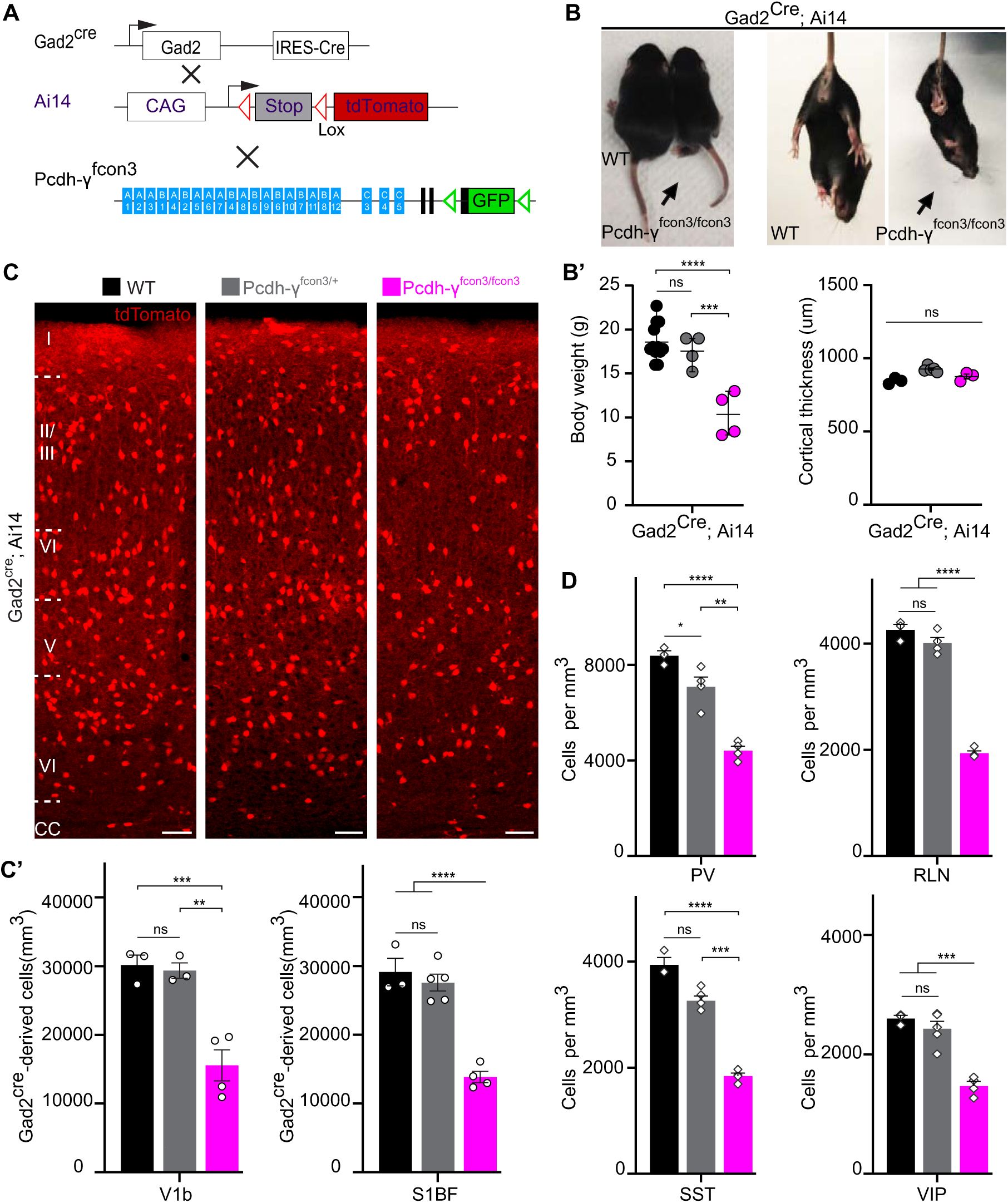
**Reduced number of GABAergic cINs in γ-Pcdh deficient mice. A**. Mutant mice with loss of Pcdh-γ in GABAergic neurons were generated by crossing conditional Pcdh-γ^fcon3^ mice to Pan-GABAergic Cre driver (Gad2) mice. The conditional Ai14 allele was used to fluorescently label Gad2-expressing cells. **B, B’.** Photographs of P21 Gad2^cre^;Ai14 Pcdh-γ^+/+^ (Pcdh-γ WT, left) and Gad2^cre^;Ai14 Pcdh-γ^fcon3/fcon3^ (Pcdh-γ mutant, right) mice. **B’**, Body weight and cortical thickness measurements in P30 Gad2^cre^;Ai14 Pcdh-γ^+/+^ (Pcdh-γ WT), Gad2^cre^;Ai14 Pcdh-γ^fcon3/+^ (Pcdh-γ HET), and Gad2^cre^;Ai14 Pcdh-γ^fcon3/fcon3^ (Pcdh-γ mutant) mice. One-way ANOVA, F=23.04, ***p=0.0003,****p<0.0001; n= 12 Pcdh-γ WT mice, n= 4 Pcdh-γ HET mice and n= 4 Pcdh-γ mutant mice. **C, C’.** Photographs of coronal sections in primary visual cortex (V1b) of P30 Gad2^cre^;Ai14 Pcdh-γ WT(left), Pcdh-γ HET (middle) and Pcdh-γ mutant (right) mice. Scale bar, 100um. **C’**, Quantifications of tdTomato+ cell density in V1b and somatosensory (S1BF) cortex of P30 Gad2^cre^; Ai14 Pcdh-γ WT(black), Pcdh-γ HET (grey) and Pcdh-γ mutant (magenta) mice. One-way Anova; F=21.15 [V1b], F=39.79[S1BF], **p=0.0027, ***p=0.0019, ****p<0.0001; n= 3 [V1b and S1BF] Pcdh-γ WT; n= 4 [V1b], n=5 [S1BF] Pcdh-γ HET; n=4 [V1b and S1BF] Pcdh-γ mutant Gad2^cre^;Ai14 mice. **D.** Quantifications of cIN subtype density in V1b cortex at P30. All four non-overlapping cIN subtypes, PV, SST, RLN and VIP, were similarly reduced in numbers in Gad2^cre^;Ai14; Pcdh-γ^fcon3/fcon3^ mice (Pcdh-γ mutant, magenta) compared to controls (black and grey). One-way ANOVA; F=41.47 [PV], F=44.80 [SST], F=206.6 [RLN], F =32.03 [VIP]; *p=0.0465, **p=0.0003,***p=0.0002, ****p<0.0001.

**Figure 3.**
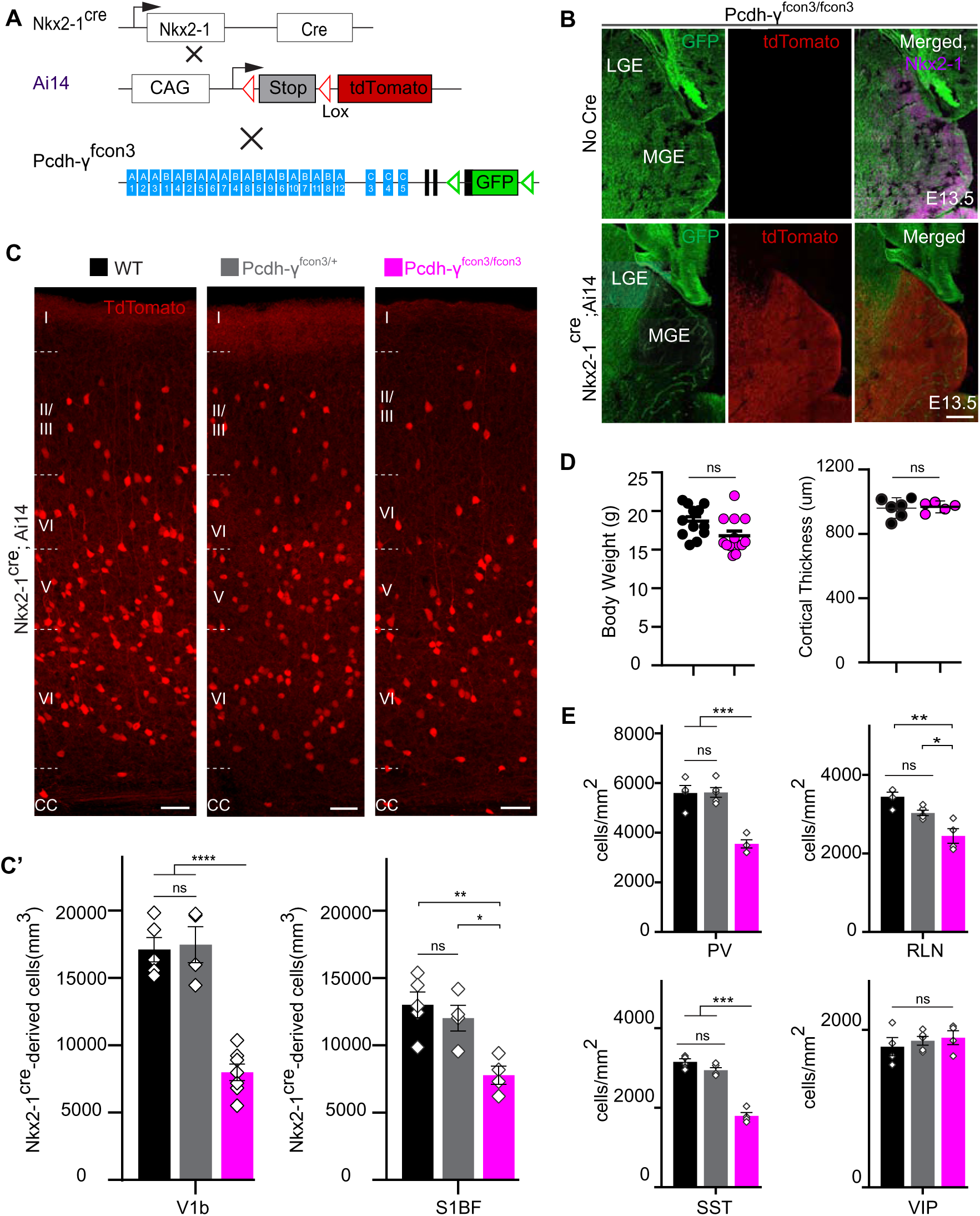
Loss of γ-Pcdh genes targeted to Nkx2-1 expressing cells results in selective loss of cIN derived from the MGE. A. Mutant mice with loss of Pcdh-γ in MGE-derived cIN were generated by crossing Pcdh-γ^fcon3^ mice to the the Nkx2-1^cre^ mouse line. The conditional Ai14 allele was used to fluorescently label MGE-derived cells. **B.** Pcdh-γ^fcon3/fcon3^ mice (top panels) carrying the Pcdh-γ mutant allele, but not Cre, show robust expression of GFP in the MGE. In contrast, in Nkx2-1^Cre^; Ai14 Pcdh-γ^fcon3/fcon3^ mice, carrying the Pcdh-γ mutant allele and expressing Cre (lower panels), GFP expression was eliminated from the MGE. The few cells left expressing GFP in the MGE are blood vessels and Ai14 negative. **C, C’**. Photographs of coronal sections of the binocular region of the primary visual cortex (V1b) in Nkx2-1^Cre^; Ai14 Pcdh-γ^+/+^ (Pcdh-γ WT), Nkx2-1^Cre^;Ai14 Pcdh-γ^fcon3/+^ (Pcdh-γ HET) and Nkx2-1^Cre^;Ai14 Pcdh-γ^fcon3/fcon3^ (Pcdh-γ mutant). Scale bar, 100um. C’ Quantification of the density of Ai14+ cells in V1b and S1BF cortex of P30 Nkx2-1^Cre^;Ai14 Pcdh-γ WT (black), Pcdh-γ HET (grey) and Pcdh-γ mutant (magenta) mice. The number of Nkx2.1-derived cells was significantly reduced in Nkx2-1^Cre^;Ai14 Pcdh-γ mutant mice compared to controls. ANOVA; F=40.92 [V1b], F= 9.151 [S1BF], *p= 0.0277, **p=0.0040, ****p<0.0001; n= 5 [V1b, S1BF] Pcdh-γ WT, n= 4 [V1b], 5 [S1BF] Pcdh-γ HET, n= 7 [V1b], 4 [S1BF] Pcdh-γ mutant Nkx2-1^Cre^;Ai14 mice. **D.** Body weight and cortex thickness measurements in Nkx2-1^Cre^;Ai14 Pcdh-γ WT (black) and Pcdh-γ mutant (magenta) mice at P30. Body weight and cortical thickness were not significantly affected by loss of Pcdh-γ. ANOVA, F= 3.642 [weight], F=1.479 [cortical thickness]; n= 12 [weight], n= 4 [cortical thickness] Pcdh-γ WT mice; n= 7 [weight]; n=13 [weight] and n= 5 [cortical thickness] Pcdh-γ mutant Nkx2-1^Cre^;Ai14 mice. Quantification of Ai14+ cIN subtypes in V1b mouse cortex at P30. Nkx2-1^Cre^;Ai14 Pcdh-γ mutant mice (magenta) had significantly reduced numbers of MGE-derived parvalbumin (PV)+, somatostatin (SST)+ and Rellin (RLN)+ cells compared to controls (black & grey). In contrast VIP+ cells, which are derived from the CGE, were not significantly affected. ANOVA; F = 26.95 [PV], F = 89.96 [SST], F = 14.82 [RLN], F = 0.4266 [VIP]; *p = 0.0182, **p = 0.0008, ***p < 0.0003, ****p < 0.0001; n = 4 Pcdh-γ WT, n = 5 Pcdh-γ HET and n = 4 Pcdh-γ mutant mice per subtype of cIN analyzed.

However, the density of Ai14 positive cells in somatosensory and visual cortex was roughly halved in homozygous *Gad2^cre^; Ai14; Pcdh-γ^fcon3/fcon3^* animals, compared to wild type and heterozygous littermates (**Figure 2C& C’**). The density of cINs stained positive for PV and SST (MGE-derived), VIP (CGE-derived) or RLN (derived from both the MGE and CGE) was significantly reduced in the visual cortex of homozygous *Gad2^cre^; Ai14; Pcdh-γ^fcon3/fcon3^* mice (**Figure 2D&Figure 2- Figure supplement 1)**. Taken together, these experiments indicate that the embryonic loss of Pcdh-γ function in GABAergic progenitor cells leads to a drastically reduced number of cINs in the neocortex, affecting all cIN subtypes similarly.

The developmental defects observed in *Gad2^cre^; Ai14; Pcdh-γ^fcon3/fcon3^* mutant mice may indirectly affect the survival of cIN in a non-cell autonomous manner. We thus decided to restrict the Pcdh-γ loss of function to MGE/POA (preoptic area) progenitors by means of the *Nkx2-1^cre^* mice (Xu, Tam and Anderson, 2008).

MGE/POA progenitors give rise to the majority of mouse cINs, including PV and SST interneurons. NKX2-1 expression is detected in the ventral telencephalon from embryonic day (E) 9.5 (Shimamura *et al*., 1995; Sandberg *et al*., 2016) and is downregulated in most cINs as they migrate into the developing neocortex (Nóbrega-Pereira *et al*., 2008). *Pcdh-γ^fcon3^* mice were crossed to *Nkx2-1^Cre^* mice. As described above, the Ai14 allele was again used to visualize the recombined cells (**Figure 3A**). Homozygous *Nkx2-1^Cre^; Ai14; Pcdh-γ^fcon3/fcon3^* embryos lost GFP expression specifically the MGE and the preoptic regions (**Figure 3B**), consistent with full recombination, and loss of Pcdh-γ function in cells derived from the Nkx2.1 lineage.

At P30, *Nkx2-1^Cre^; Ai14; Pcdh-γ^fcon3/fcon3^* mice displayed a dramatic reduction (∼50%) in the number of MGE-derived tdTomato+ cells (**Figure 3C& C’)**, both in the visual and somatosensory cortex. MGE-derived PV and SST interneuron number was similarly greatly reduced in these animals. However CGE-derived VIP interneuron density was similar to that of control animals (**Figure 3E&Figure 3- Figure supplement 1)**. A smaller, but significant reduction in the RLN positive cIN population was observed, in agreement with the notion that a subpopulation of RLN cells are born in the MGE (Miyoshi *et al*., 2010). Consistently, layer 1 RLN+ cells, which are largely derived from the CGE (Miyoshi *et al*., 2010), were not affected by Pcdh-γ loss of function, but RNL cells in deeper layers 2-6 (which many are MGE-derived and also positive for SST) showed reduced numbers (**Figure 3- Figure supplement 2)**. Together these results show that embryonic loss of Pcdh-γ function in Nkx2-1-positive progenitors results in a significant reduction in the number of MGE/POA-derived cINs.

### Pcdh-γ function is not required for the proliferation and migration of cIN precursors

The reduction in the number of cINs in *Nkx2-1^Cre^; Ai14; Pcdh-^fcon3/fcon3^* mice was not a result of abnormal cortical thickness or abnormal layer distribution, as these measures were similar across genotypes in P30 mice (**Figure 3C**). We next asked whether migration or proliferation defects in the cIN progenitor population could lead to a reduced cIN density in Pcdh-γ mutant mice. Quantification of the number of dividing cells in the ventricular or subventricular zones at E13.5 and E15.5, using the mitotic marker Phosphohistone H3 (PH3), showed no differences in the number of mitotic cells in the MGE between *Nkx2-1^Cre^;Ai14; Pcdh-γ^fcon3/fcon3^* mice and controls (**Figure 4A& B**). Migration of young cINs into cortex was also not affected in *Nkx2-1^Cre^; Ai14; Pcdh-γ^fcon3/fcon3^*. The tdTomato+ cells in the cortex displayed a similar migratory morphology in *Nkx2-1^Cre^; Ai14; Pcdh-γ^fcon3/fcon3^* embryos and controls. Consistently, the number of migrating cells in cortex in the marginal zone (MZ), the subplate (SP), and the intermediate and subventricular zone (IZ/SVZ) was equivalent between Pcdh-γ mutant embryos and controls at E15.5 (**Figure 4C& D**). These findings indicate that loss of Pcdh-γ did not affect the proliferation of MGE progenitors or the migration of young MGE-derived cINs into the developing neocortex.

**Figure 4.**
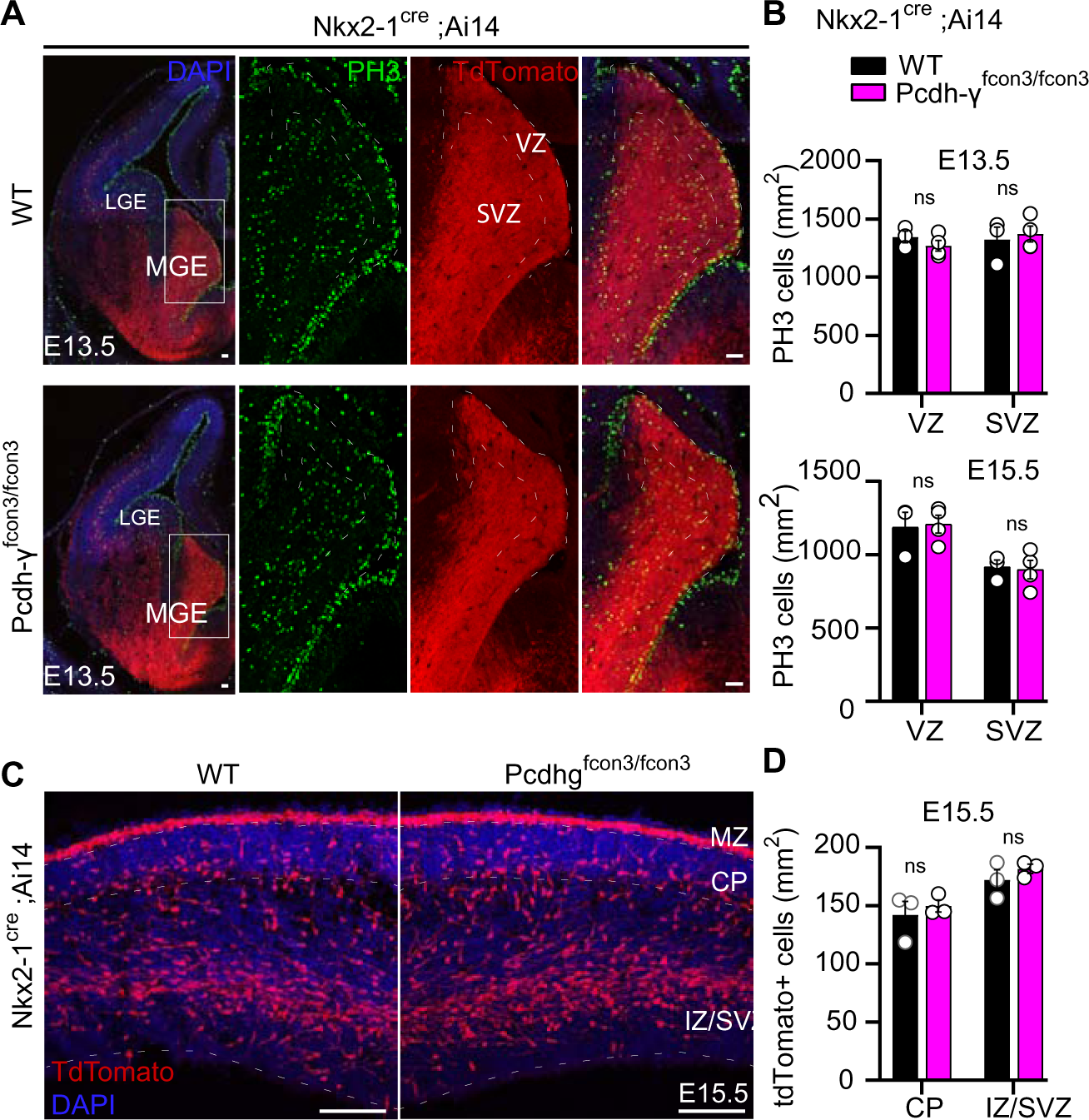
Proliferation and migration are not affected by the loss of γ-Pcdhs in Nkx2-1 expressing cells. A. Photographs of coronal sections through the embryonic forebrains of E13.5 Nkx2-1^Cre^;Ai14Pcdh-γ^+/+^ (Pcdh-γ WT, top panels) or Nkx2-1^Cre^;Ai14;Pcdh-γ^fcon3/fcon3^ (Pcdh-γ mutant, bottom panels). Close-up photographs of the MGE from Nkx2-1^Cre^;Ai14 Pcdh-γ WT (right panels) or Pcdh-γ mutant (bottom, right panels) embryos. Robust reporter activity (Ai14) was observed in the MGE. Dividing cells were labeled using the mitotic marker PH3. Note that the size of the MGE and number of PH3 proliferating cells was similar in the mutant and control brains. Scale bars, 50um **B.** Quantification of PH3 positive cells from MGE ventricular (VZ) and subventricular zone (SVZ) in E13.5 (top) and E15.5 (bottom) Nkx2-1^Cre^;Ai14 Pcdh-γ WT (black bars) and Pcdh-γ mutant (magenta bars) embryos. **C.** Photographs of coronal sections of dorsal cortex at E15.5 showing the migrating MGE-derived cIN in Nkx2-1^Cre^;Ai14 Pcdh-γ WT (left) and Pcdh-γ mutant (right) embryos. Note the robust migratory streams of young neurons in the SVZ and in the marginal zone(MZ). From these regions cells disperse into the intermediate zone (IZ) and cortical plate (CP). Similar number of migrating cIN were observed in mutants and controls. Scale bar, 100um. **D.** Quantifications of number of migrating MGE-derived cINs in the CP and in the IZ/SVZ of Nkx2-1^Cre^;Ai14 Pcdh-γ WT (black) and Pcdh-γ mutant (magenta) mice. No significant differences were detected in the number of Ai14 + migrating cells in Pcdh-γ mutant and WT controls. For **B** and **D**, Two-tailed Student’s unpaired t-test; For PH3 quantification, p= 0.3557 [VZ, E13.5], p= 0.7026 [SVZ, E13.5], p= 0.7000[ VZ, E15.5], p=0.4548 [SVZ, E15.5]; n=3 [VZ, E13.5 & 15.5,] Pcdh-γ WT mice and n=4 [SVZ, E13.5], n=3 [SVZ, E15.5] Pcdh-γ mutant mice. For quantification of migrating cIN, p= 0.3973 [E15.5, IZ-SVZ], p=0.5785 [E15.5 CP].

### Accentuated cIN cell death in Pcdh-γ mutants

A wave of programmed cell death eliminates ∼40% of the young cINs shortly after their arrival in the cortex (Southwell *et al*., 2012; Wong *et al*., 2018). This wave starts at ∼P0, peaks at P7 and ends at ∼P15. We next asked if the reduced cIN density observed in Pcdh-γ mutant mice could stem from a heightened number of mutant cINs undergoing apoptosis. Such cells were immunolabeled using an antibody directed against cleaved-Caspase 3 (cc3). Since cc3 positive cells are relatively rare, our analysis was performed throughout the entire neocortex, at P0, 3, 7, 10 and 15. Similarly to their wild type littermates, *Nkx2-1^Cre^;Ai14; Pcdh-γ^fcon3/fcon3^* homozygous mice displayed a wave of programmed cell death peaking at P7 (**Figure 5A& B**).

**Figure 5.**
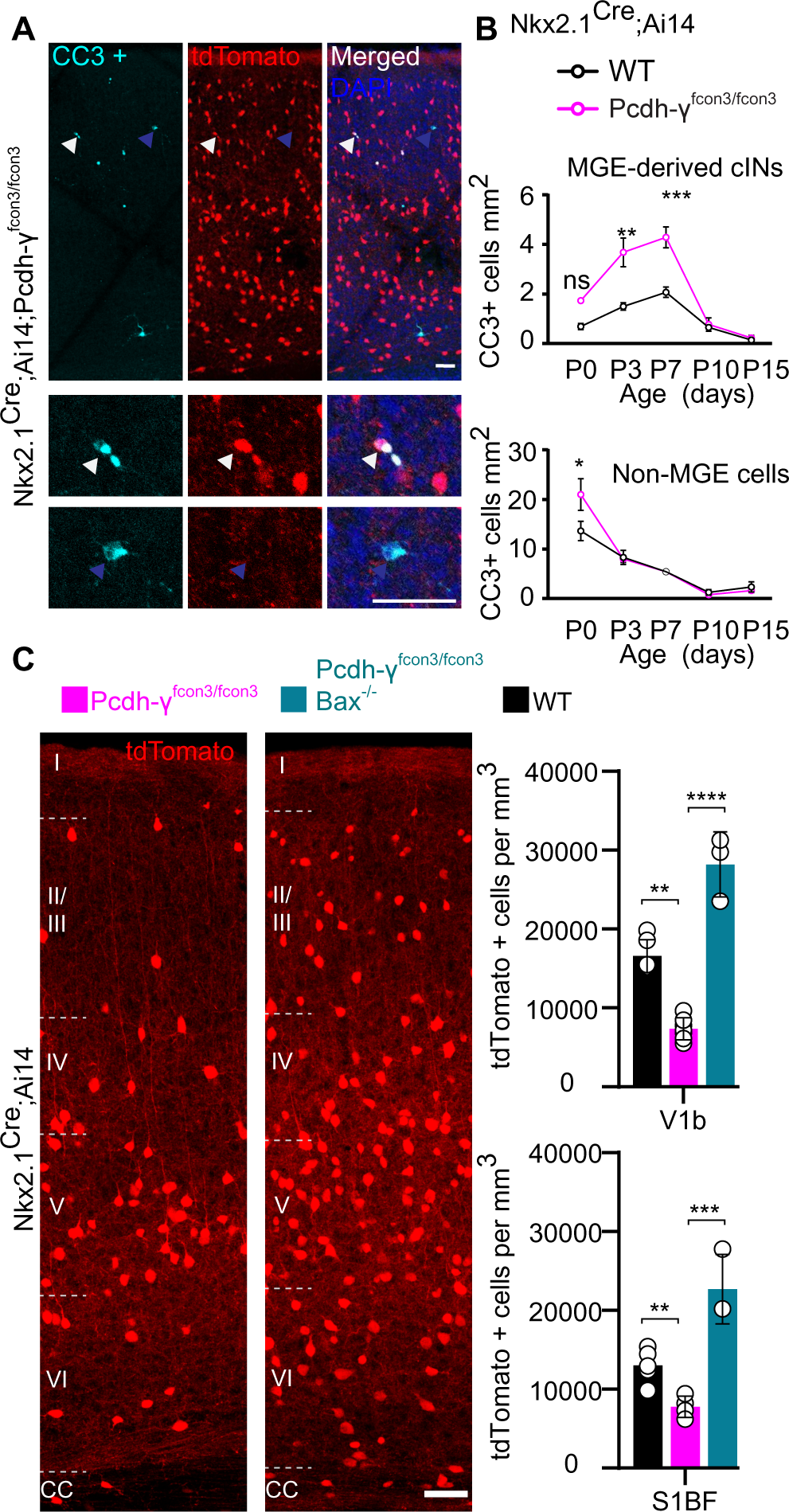
Increased programmed cell death in Pcdh-γ mutants is rescued in Pcdh-γ-bax null animals. A. Photographs of coronal sections through a Nkx2-1^Cre^;Ai14 Pcdh-γ^fcon3/fcon3^ (Pcdh-γ mutant) P7 mouse cortex (top), showing Ai14+ cIN and cleaved caspase 3 positive cells (CC3+). Close-up photographs (bottom) of Ai14+, CC3+ (white Arrowheads) and Ai14-, CC3+ (blue Arrowheads) cells. **B.** Quantification of the density of Ai14+,CC3+ (MGE-derived, top graph) cells from Nkx2-1^Cre^;Ai14 Pcdh-γ WT (black line) and Pcdh-γ mutant (magenta line) mice. Quantification of the density of Ai14-,CC3+ (non-MGE-derived, bottom graph) cells from Nkx2-1^Cre^;Ai14 Pcdh-γ WT (black line) and Pcdh-γ mutant (magenta line) mice. Note that the number of CC3+ cells was significantly increased in the MGE-derived population in Pcdh-γ mutant mice, and coincides with the normal period of programmed cell death for cIN in WT mice. Two-way ANOVA, F = 16.83, **p = 0.0029 [P3], **p = 0.0044 [P7]; n= 4 [P0 and P10], n = 3 [P3, P7 and P15] Pcdh-γ WT mice; n= 3 [P0,P10 and P15], n = 7 [P3] and n=5 [p7]Pcdh-γ mutant mice.. **C.** Coronal sections through the binocular region of the primary visual cortex (V1b) of Nkx2-1^Cre^;Ai14 (Pcdh-γ WT, left) and Nkx2-1^Cre^;Ai14;Pcdh-γ^fcon3/fcon3^; Bax^-/-^(Pcdh-γ mutant, Bax null, right) mice at P30. Quantifications of the density of cINs in the V1b (top) and S1BF (bottom). Note that genetic removal of Bax in Pcdh-γ mutant mice rescues cell death; both the endogenous normal cIN elimination and the additional cell death observed in Pcdh-γ mutant animals. ANOVA, P value = 0.0001 [V1b], P value <0.0001 [S1BF]; **p = 0.0037,, ***p = 0.0022,****p = 0.0012; n = 8 [V1b] and n = 6 [S1BF] Pcdh-γ WT mice; n = 12[V1b] and n = 5 [S1BF] Pcdh-γ mutant mice; n = 3 [V1b] and n = 3 [S1BF] Pcdh-γ mutant; Bax null mice.

However, Pcdh-γ mutant mice had significantly higher numbers of tdTomato+/cc3+ cells compared to controls. We also examined the proportion of cc3+ cells that were tdTomato negative (un-recombined cells that would notably include pyramidal cells, CGE-derived cINs, and glial cells). With the exception of a small, but significant increase observed at P0, we found no significant difference in the number of cc3+/tdTomato-cells between genotypes (**Figure 5B, bottom graph)**. This suggests that the survival of neighboring Pcdh-γ-expressing cells is not impacted by the loss of Pcdh-γ-deficient MGE/POA-derived cINs. Importantly, the homozygous deletion of the pro-apoptotic Bcl-2-associated X protein (BAX) rescued cIN density in the Pcdh-γ mutant mice to levels similar to those observed in *BAX^-/-^* mice (Southwell *et al*., 2012) (**Figure 5C**). The above results indicate that loss of Pcdh-γ in MGE/POA-derived cIN enhances their culling through programmed cell death during the developmental period when these cells are normally eliminated.

### Loss of Pcdh-γ does not affect survival of cINs after the period of programmed cell death

We then asked whether Pcdh-γ expression is also required for the survival of cINs past the period of programmed cell death. To address this question we took advantage of the *PV^Cre^* transgene (Hippenmeyer *et al*., 2005) that becomes activated specifically in PV interneurons starting at around ∼P16 (**Figure 6&Figure 6- Figure supplement 1**). Quantifications of tdTomato+ cell density in *PV^Cre^; Ai14; Pcdh-γ^fcon3/fcon3^* and PV-cre; Ai14 mice at P60-P100 revealed no significant differences between homozygous and control mice (V1b and S1BF) (**Figure 6D& E**). The SST-Cre line that shows recombination activity specifically in SST interneurons at embryonic stages. Using this allele, we could demonstrate that, as observed above using the Nkx2-1-Cre line, loss of Pcdh-γ in developing SST interneurons led to a reduction in the cortical density for these cells at P30 (**Figure 6A-C**). Together, our results demonstrate that Pcdh-γ loss of function reduces survival, specifically during the endogenous period of cortical interneuron cell death resulting in reduced cortical density of cINs.

**Figure 6.**
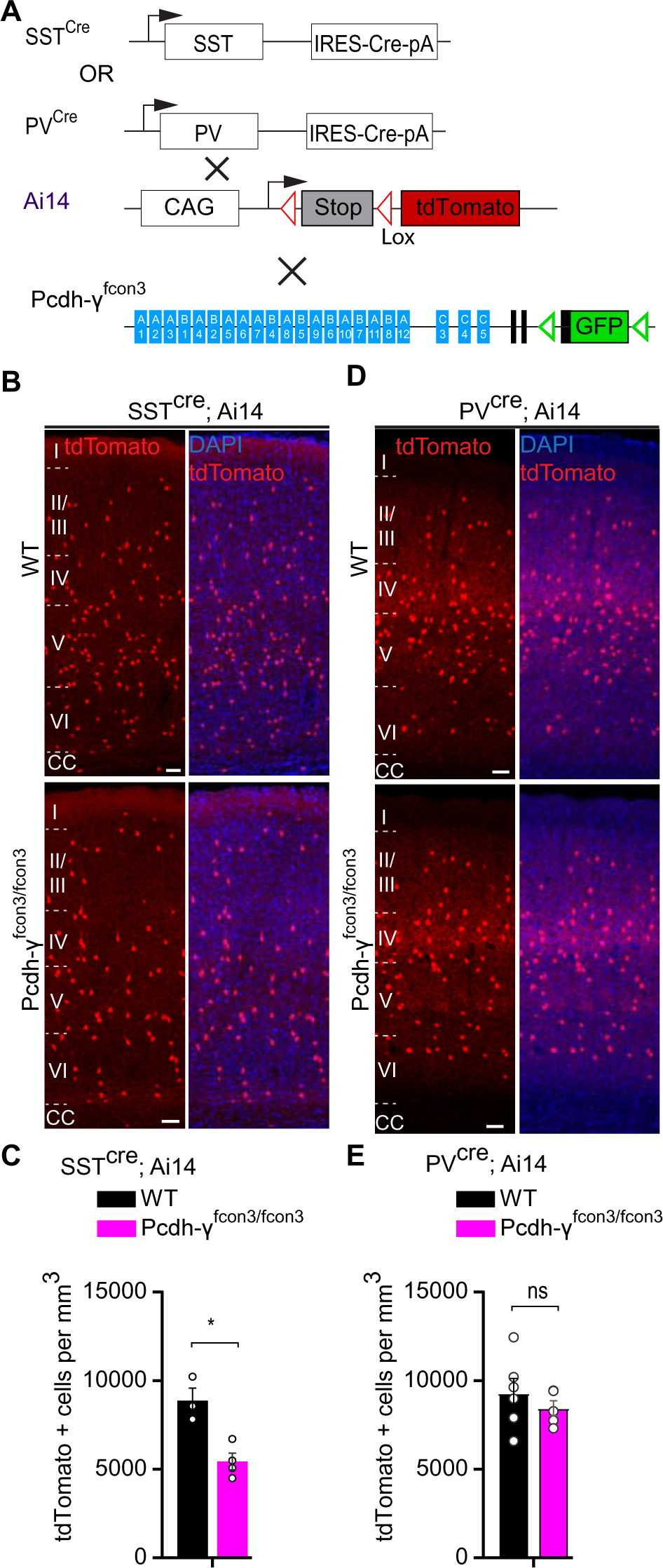
Pcdh-γ function is not required for the survival of PV cINs after the period of programmed cell death. A. Mutant mice with loss of Pcdh-γ in SST or PV cells were generated by crossing conditional Pcdh-γ^fcon3^ mice to mice carrying Cre under SST or PV. The conditional Ai14 allele was used to fluorescently label SST or PV cells. **B.** Photographs of coronal sections of the binocular region of the primary visual cortex (V1b) of P30 SST^Cre^; Ai14 Pcdh-γ^+/+^ (Pcdh-γ WT, top left) and SST^Cre^; Ai14 Pcdh-γ^fcon3/fcon3^ (Pcdh-γ mutant, bottom left) mice. Scale bars, 50um. **C.** Quantifications of the density of Ai14+ cIN in V1b cortex of Pcdh-γ WT (black) and Pcdh-γ mutant (magenta) SST^Cre^; Ai14 P30 mice. Two-tailed Student’s unpaired t-test, **p= 0.0084, n=5 mice of each genotype. **D.** Coronal sections through the primary visual cortex (V1b) in PV^Cre^; Ai14 Pcdh-γ^+/+^ (Pcdh-γ WT, top right) and PV^Cre^; Ai14;Pcdh-γ^fcon3/fcon3^ (Pcdh-γ mutant, bottom right) mice at P30. Scale bars, 50um. **E.** Quantifications of the density of Ai14+ cIN in V1b cortex of Pcdh-γ WT and Pcdh-γ mutant PV^Cre^; Ai14 mice at P60-100. Two-tailed Student’s unpaired t-test, p = 0.4275, n= 5 mice of each genotype.

### Alpha and beta Pcdhs do not impact cIN survival

Previous studies indicate that Pcdhs form tetrameric units that include members of the α-, β-, and *γ*-clusters (Dietmar Schreiner, 2010; Thu *et al*., 2014). We, therefore asked whether α and β Pcdhs also contributed to cIN cell death. Mice that carry a conditional deletion of the entire α cluster (*Pcdh-α^acon/acon^*) were crossed to the *Nkx2-1^Cre^; Ai14* line, resulting in removal of the Pcdh-α cluster genes, specifically from MGE/POA progenitor cells (**Figure 7A**). *Nkx2-1^Cre^; Ai14; Pcdh-α^acon/acon^* mice were viable, fertile, and displayed normal weight (**Figure 7B, top graph**). We observed that cIN density in the visual cortex of *Nkx2-1^Cre^; Ai14; Pcdh-α^acon/aco^*^n^ mice at P30 was similar to that of *Nkx2-1^Cre^; Ai14* mice (**Figure 7B**). To determine if the Pcdh-β cluster affected MGE/POA-derived cIN survival, constitutive Pcdh-β cluster knockout mice were crossed to *Nkx2-1^Cre^; Ai14* mice (**Figure 7a**). Similarly to the deletion of Pcdh-α cluster, mice carrying a deletion of the entire Pcdh-β cluster, are viable, fertile and of a normal weight (**Figure C, top graph**) (Chen *et al*., 2017) The density of cIN was similar between mice lacking β-Pcdhs and controls (**Figure 7C**). The above results indicate that unlike the Pcdh-γ cluster that is essential for the regulation of cIN elimination, the function of α- or β-Pcdhs is dispensable for the survival of MGE/POA-derived cINs.

**Figure 7.**
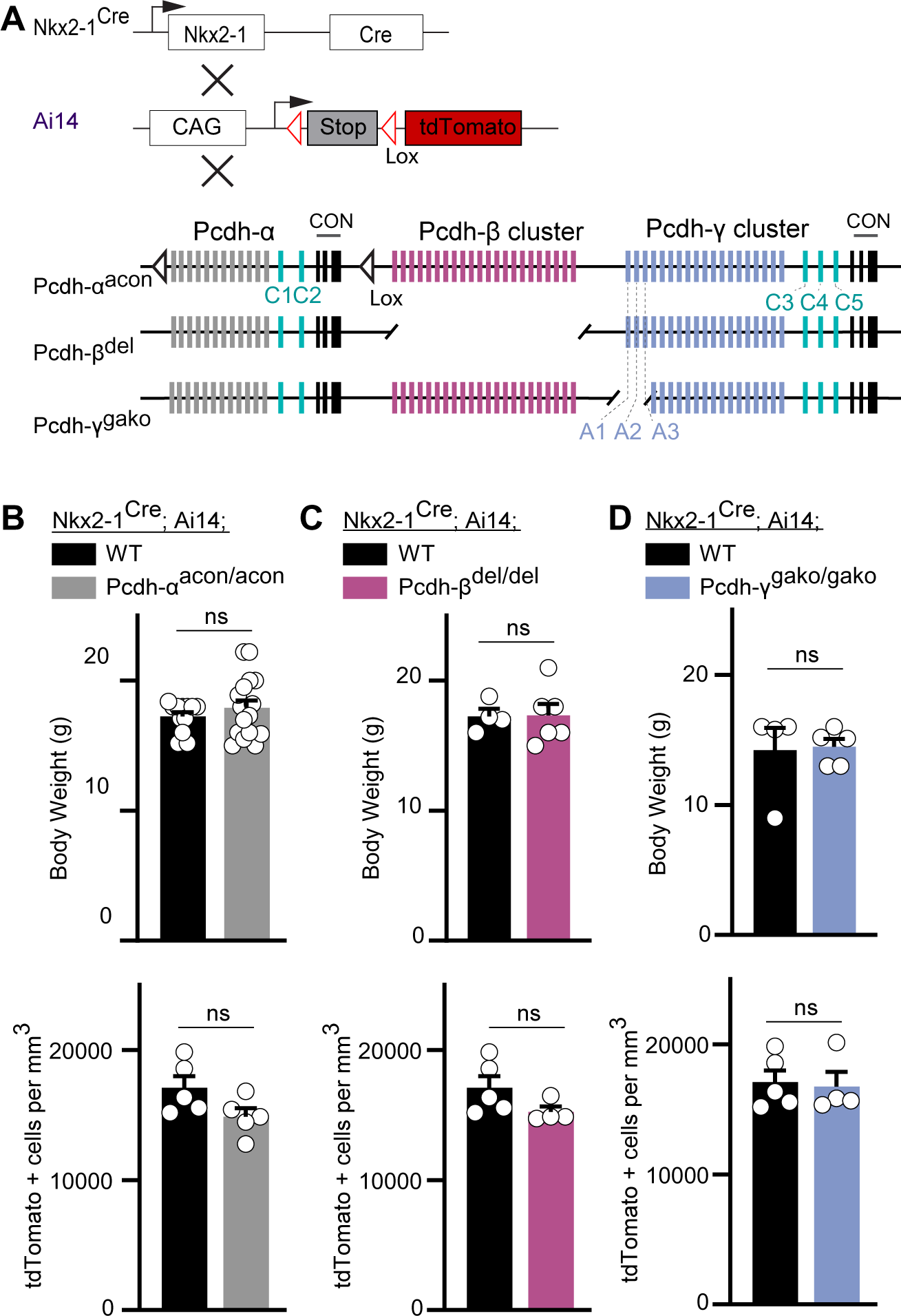
Loss of Pcdh-α, Pcdh-β, or Pcdh-γ A1-A2-A3 genes does not affect the survival of MGE-derived cIN. A. Mutant mice with loss of Pcdh-α, Pcdh-β or Pcdh-γ A1-A2-A3 genes in MGE-derived cIN were generated by crossing Pcdh-γ^acon^, Pcdh-β^del^, or Pcdh-γ^gako^ mice to the the Nkx2-1^Cre^ mouse line. The conditional Ai14 allele was used to fluorescently label MGE-derived cells. **B.** (Top) Measurements of body weight in P30 Nkx2-1^Cre^;Ai14;Pcdh-α^+/+^ (Pcdh-α WT, black) and Nkx2-1^Cre^;Ai14;Pcdh-α^acon/acon^ (Pcdh-α mutant, grey) mice. (Bottom) Quantification of the density of MGE-derived cIN in the primary visual cortex (V1b) of Pcdh-α WT (black) and Pcdh-α mutant (grey) Nkx2-1^Cre^;Ai14 P30 mice. **C.** (Top) Measurements of body weight in P30 Nkx2-1^Cre^;Ai14;Pcdh-β^+/+^ (Pcdh-β WT, black) and Nkx2-1^Cre^;Ai14;Pcdh-β^del/del^ (Pcdh-β mutant, pink) mice. (Bottom) Quantification of the density of MGE-derived cIN in the primary visual cortex (V1b) of Pcdh-β WT (black) and Pcdh-β mutant (pink) Nkx2-1^Cre^;Ai14 P30 mice. **D.** (Top) Measurements of body weight in Nkx2-1^Cre^;Ai14;Pcdh-γ^+/+^ (Pcdh-γ WT, black) and Nkx2-1^Cre^;Ai14;Pcdh-γ^gako/gako^(Pcdh-γ A1-A2-A3 mutant, blue) P30 mice. (Bottom) Quantification of the density of MGE-derived cIN in the primary visual cortex (V1b) of Pcdh-γ WT(black) and Pcdh-γ A1-A2-A3 mutant (blue) Nkx2-1^Cre^;Ai14 P30 mice. Two-tailed Student’s unpaired t-test [B-D]; n= 12[weight], n=5 [V1b cell density] Pcdh-α WT Nkx2-1^Cre^;Ai14 mice; n= 17[weight], n=5 [V1b cell density] Pcdh-α mutant Nkx2-1^Cre^;Ai14 mice; n= 4[weight], n= 5[V1b cell density] Pcdh-β WT Nkx2-1^Cre^;Ai14 mice; n= 5[weight], n= 4[V1b cell density] Pcdh-β mutant Nkx2-1^Cre^;Ai14 mice; n= 4[weight],n= 5[V1b cell density] Pcdh-γ WT Nkx2-1^Cre^;Ai14 mice; n= 5[weight],n= 4[V1b cell density] Pcdh-γ A1-A2-A3 mutant Nkx2-1^Cre^;Ai14 mice.

### Loss of Pcdh-γ does not affect cIN dispersion after transplantation but affects their survival

In order to compare the timing and extent of migration, survival, and maturation of cINs of different genotypes within the same environment, we co-transplanted into the cortex of host animals, MGE-derived cIN precursor cells expressing red and green fluorescent proteins. MGE cIN precursors were either derived from E13.5 Gad67-GFP embryos (Pcdh-γ WT controls) or from *Nkx2-1^Cre^;Ai14* embryos that were either Pcdh-γ WT or Pcdh-γ mutant (**Figure 8A**). We first confirmed that MGE cells WT for Pcdh-γ, but carrying the two different fluorescent reporters displayed no differences in their survival. Equal numbers of Gad67-GFP cells (Pcdh-γ WT GFP+) and *Nkx2-1^Cre^; Ai14* cells (Pcdh-γ WT Ai14+) were co-transplanted into the neocortex of neonatal recipients. The proportion of surviving GFP+ and tdTomato+ cells at 3, 6, 13 and 21 days after transplantation (DAT) was measured (**Figure 8A& B, top graph**). The contribution of each cell population to the overall pool of surviving cells was found to be ∼ 50% at 3 DAT, and remained constant at 6, 13 and 21 DAT (**Figure 8B**, **top graph**). This experiment indicates that the fluorescent reporters (GFP or tdTomato) or breeding background does not affect the survival of MGE cINs in this assay. Next, we co-transplanted equal numbers of Gad67-GFP cells (Pcdh-γ WT) and *Nkx2-1^Cre^; Ai14; Pcdh-γ^fcon3/fcon3^* cells (Pcdh-γ mutant) into the cortex of WT neonatal recipients. As above, we measured the proportion of surviving GFP+ and tdTomato+ cells at 3, 6, 13 and 21 DAT (**Figure 8A**). Similar numbers of GFP+ and tdTomato+ cells were observed at 3 and 6 DAT. However, the survival fraction of the tdTomato+ population dramatically decreased at 13 and 21 DAT (**Figure 8B, bottom graph, & 8C**). Interestingly, the observed decrease in cell number in the tdTomato+ Pcdh-γ mutant population occurred when the transplanted cells reached a cellular age equivalent to that of endogenous cIN during the normal wave of programmed cell death (6DAT roughly equivalent to P0, 21DAT roughly equivalent to P15) (Southwell *et al*., 2012).

**Figure 8.**
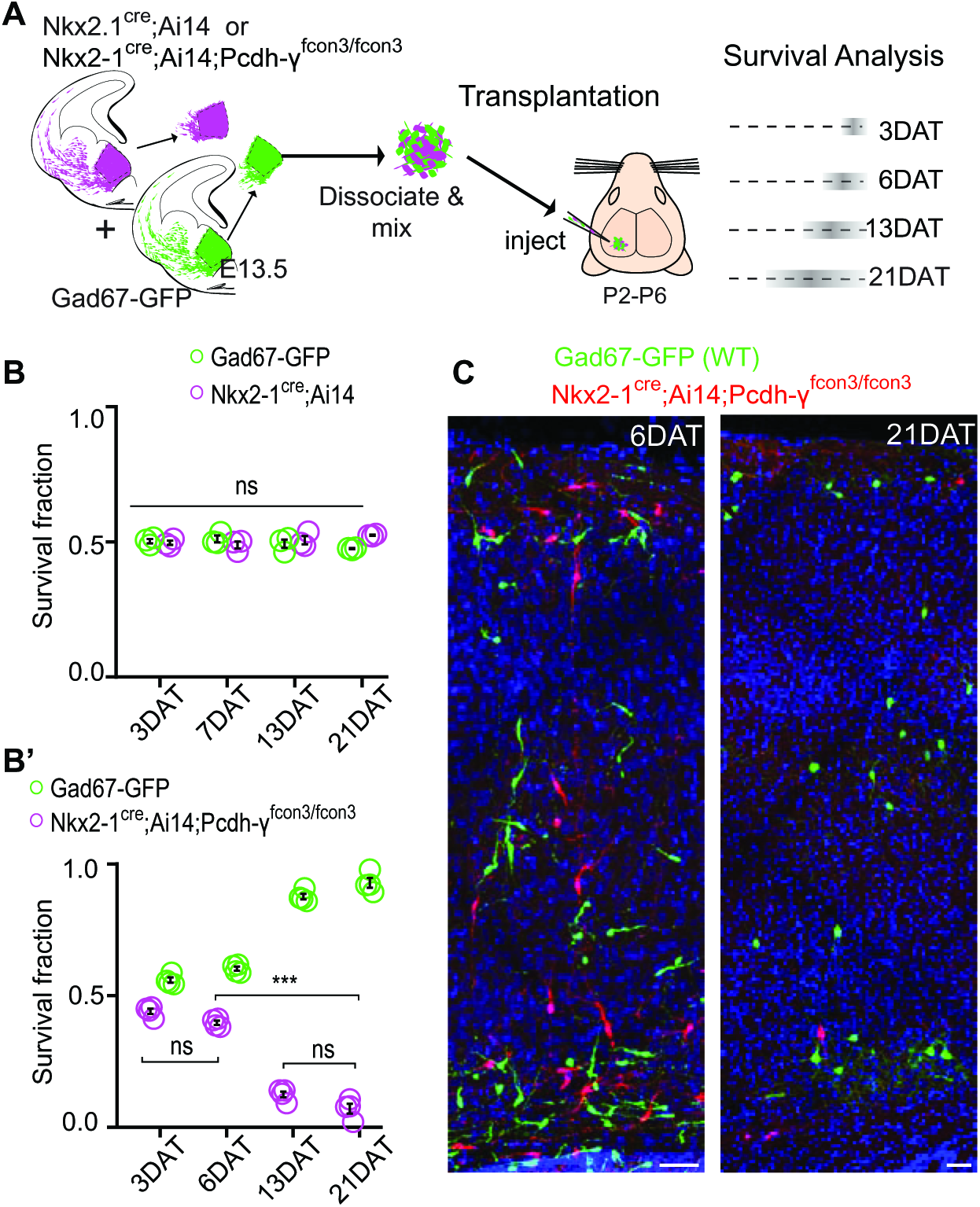
Pcdh-γ are required cIN survival after transplantation. A. Schematic of co-transplanstation of MGE-derived cIN precursors. MGE cells were derived from Nkx2-1^Cre^;Ai14 Pcdh-γ^+/+^ (Pcdh-γ WT) or Nkx2-1^Cre^;Ai14 Pcdh-γ^fcon3/fcon3^ (Pcdh-γ mutant) embryos. These cells were mixed in equal proportions with MGE cells from Gad67-GFP embryos (Pcdh-γ WT, green) and transplanted into host recipients. Cell survival was analyzed before (3DAT) and throughout the period of cell death (6-21DAT). **B,B’**. Survival fraction of co-transplanted MGE-derived cIN precursors. **B.** MGE cells were derived from Gad67-GFP (green) and Nkx2-1^Cre^;Ai14 (magenta) embryos; both GFP+ and Ai14+ cells were WT for Pcdh-γ. In this control experiment the survival was similar for both genotypes carrying the different fluorescent reporters. **B’.** MGE cells were derived from Gad67-GFP Pcdh-γ WT (green) and Nkx2-1^Cre^;Ai14 Pcdh-γ mutant (magenta) embryos. GFP+ and Ai14+ cells showed dramatic differences in their survival; the majority of cells carrying the Pcdh-γ mutant allele(magenta) were eliminated between 6 and 21 DAT. The quantifications were done at 3, 6, 13 and 21 DAT and are represented as fraction of GFP+ or Ai14+ from total cells (GFP + Ai14) per slide. ANOVA; ***P<0.0001; p= ns; 10 sections per animal; n=3 [**B**], n=4 [**B’**] mice per time point. **C.** Representative photographs of cortical sections from transplanted host mice at 6 (left) and 21 (right) DAT. Transplanted MGE cells were derived from Gad67-GFP Pcdh-γ WT (green) and Nkx2-1Cre;Ai14 Pcdh-γ mutant (red) embryos. Scale bars, 50um.

We next determined whether the survival of either Pcdh-γ WT (GFP+) or Pcdh-γ mutant (Ai14+) population was affected by their density (**Figure 9**). At 6 DAT, WT and Pcdh-γ mutant MGE-derived cells had migrated away from the injection site establishing a bell-shaped distribution (**Figure 9B& B’**). The dispersion of developing cINs lacking Pcdh-γ was indistinguishable from that of control WT cells (**Figure B’, top graph),** consistent with our observation that Pcdh-γ expression is not required for the migration of MGE-derived cINs. Strikingly, the survival fraction at 6 DAT of control Pcdh-γ WT (GFP+) and Pcdh-γ mutant (*Nkx2-1^Cre^; Ai14; Pcdh-γ^fcon3/fcon3^*) cells at the injection site or at multiple locations anterior or posterior to the site of injection was very similar (**Figure 9B’, bottom graph)**. By 21 DAT the survival of Pcdh-γ mutant (*Nkx2-1^Cre^; Ai14; Pcdh-γ^fcon3/fcon3^*) cells was dramatically reduced, but again similarly at all locations with respect to the injection site (**Figure 9B& B’).** Since the density of cIN is very different in different locations with respect to the injection site, this indicates that the survival of control Pcdh-γ WT and Pcdh-γ mutant cIN does not depend on their density.

**Figure 9.**
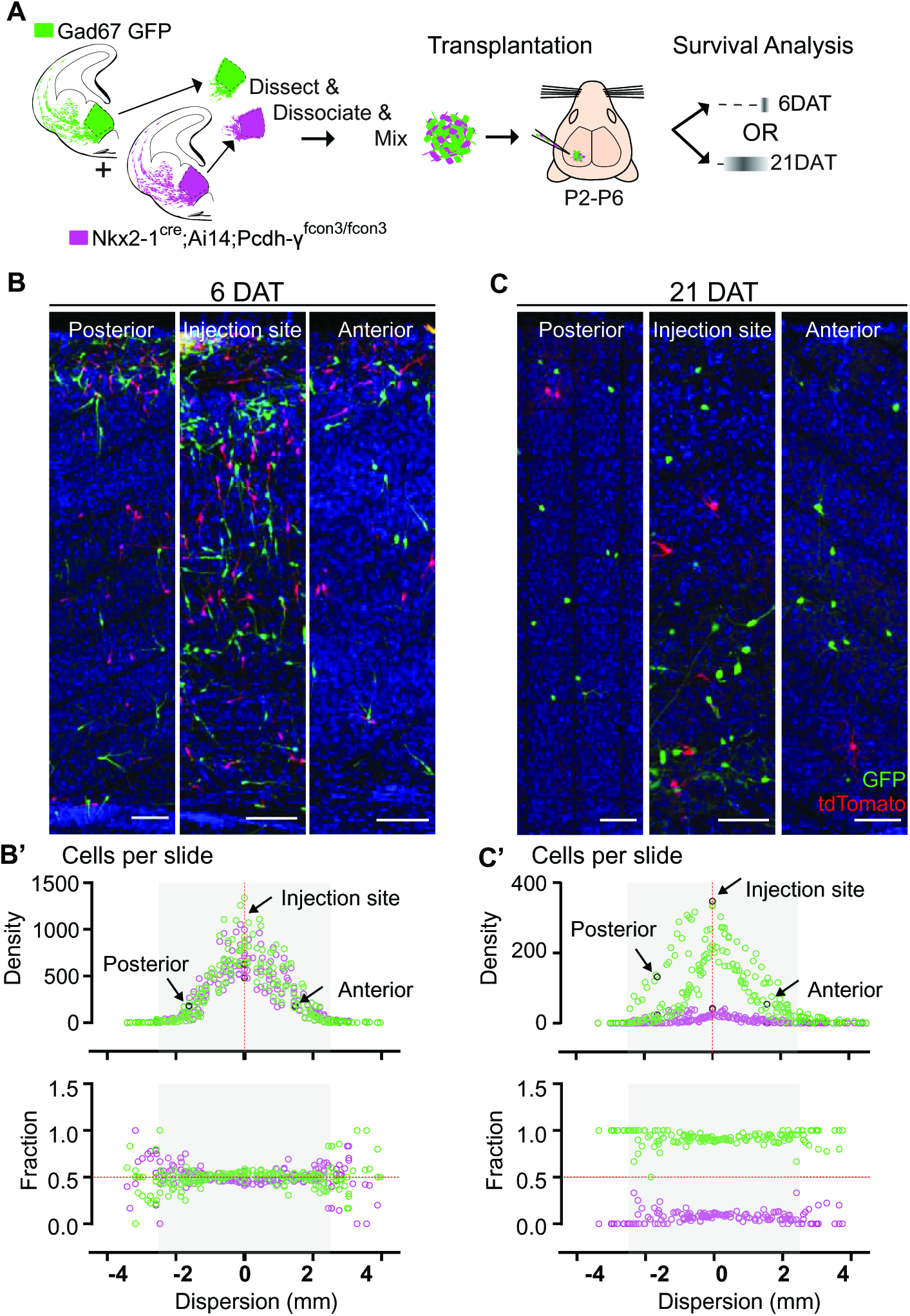
Survival of cIN, WT or mutant for Pcdh-γ, was not affected by cell density. A. MGE cells derived from Nkx2-1^Cre^;Ai14 Pcdh-γ^fcon3/fcon3^ embryos (Pcdh-γ mutant,magenta) and from Gad67-GFP embryos (Pcdh-γ WT, green) were mixed in equal numbers and transplanted into WT hosts. The survival of Ai14 and GFP labeled cIN was analyzed in every other section throughout the brain region of the transplant dispersal. **B.** Photographs of representative coronal sections at the injection site, or anterior and posterior to it, from host mice at 6DAT. Similar numbers of Ai14 and GFP labeled cIN were observed at each location. Scale bar 100um. **B’.** Dispersion analysis at 6DAT of the Pcdh-γ WT (GFP) or Pcdh-γ mutant (magenta) cells, represented as density (top) or survival fraction (bottom) as a function of distance from the site of injection in the host recipients. Note that the density of cells decreases as one moves anteriorly or posteriorly with respect to the injection site. At 6 DAT, the dispersal and survival was similar for both WT or Pcdh-γ mutant cells. **C.** Photographs of representative coronal sections at the injection site, or anterior and posterior to it, from host mice at 21DAT. Note the dramatic reduction in the number of Pcdh-γ mutant cells (Magenta) compared to the Pcdh-γ WT cells (GFP). Scale bar 100um. **C’.** Dispersion analysis at 21DAT of the Pcdh-γ WT (GFP) or Pcdh-γ mutant (Magenta) cells, represented as density (top) or survival fraction (bottom) as a function of distance from the site of injection in the host recipients. At 21 DAT, the survival fraction for the Pcdh-γ mutant cells(Magenta) was dramatically reduced and similarly affected at different locations with respect to the injection site.

In order to determine the absolute number of cINs eliminated in our co-transplantation experiments, we transplanted 50K cells of each genotype (Pcdh-γ WT and Pcdh-γ WT mutant) into host mice (**Figure 10A**).

Our baseline for survival was established at 6 DAT before the period of cIN programmed cell death. In control experiments transplanting cIN precursors derived from Nkx2-1^Cre^; Ai14 embryos (but WT for the Pcdh-γ allele) alone, we observed that ∼40% of the transplanted cIN population was eliminated between 6 and 21 DAT (**Figure 10A-C**). Therefore transplanted MGE cINs not only undergo programmed cell death during a period defined by their intrinsic cellular age but are also eliminated in a proportion that is strikingly similar to that observed during normal development (Southwell *et al*., 2012; Wong *et al*., 2018). Given these observations, we next asked how the presence of Pcdh-γ mutant cIN affected the survival of WT cIN in the transplantation setting. We co-transplanted Gad67-GFP Pcdh-γ WT (GFP+) and Nkx2-1^Cre^; Ai14 Pcdh-γ mutant (tdTomato+) MGE cIN precursors and determined the total survival of each population at 6 and 21 DAT. At 6 DAT the total number of tdTomato+ cells throughout the cortex of recipient mice was similar to that of GFP+ cells (**Figure 10D & E**). However, between 6- and 21DAT, the total number of GFP+ cells had decreased by ∼63% (**Figure 10E, compare to Figure 10C**), which suggests that WT cells die at a higher rate when co-transplanted with Pcdh-γ mutant MGE cells. Similarly, between 6- and 21DAT, the total number of tdTomato+ cells had decreased by ∼96% (**Figure 10E**). This confirms that MGE cells lacking Pcdh-γ function are eliminated in far greater numbers compared to control MGE cells, but also suggests that the presence of *Pcdh-γ* mutant cINs within a mixed population, also affects the survival of WT cINs (**compare to Figure 8**). We conclude that Pcdh-γ loss of function has both cell-autonomous and non-cell autonomous roles in the regulation of programmed cIN death.

**Figure 10.**
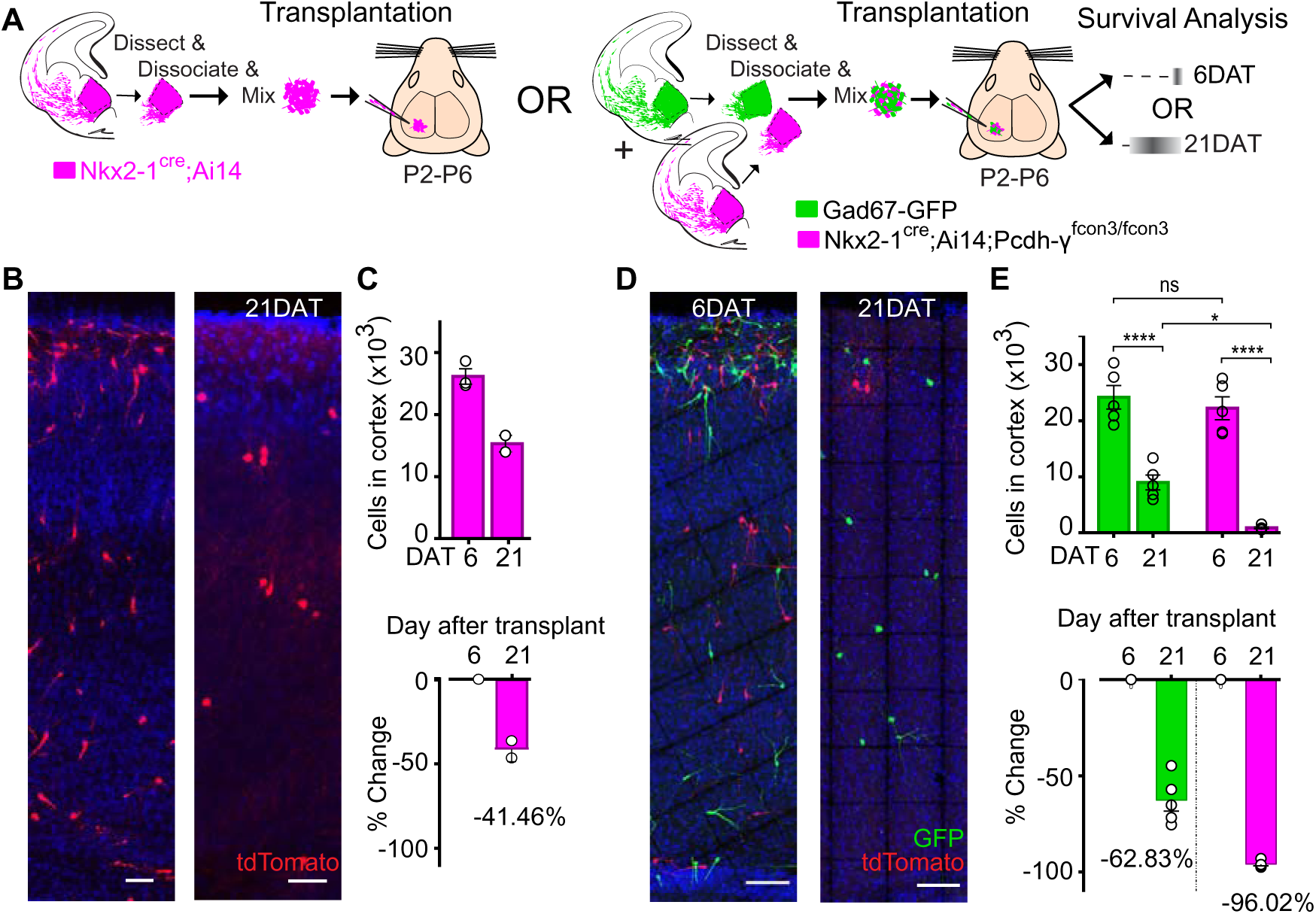
MGE cell transplantation reveals a non-cell autonomous effect of Pcdh-γ on cIN survival. A. Schematic of transplantation experiment for quantification of absolute number of transplanted MGE cells derived from Nkx2-1^Cre^;Ai14 Pcdh-γ^+/+^ alone (Pcdh-γ WT, left) OR from a mixture of equal numbers of Nkx2-1^Cre^;Ai14 Pcdh-γ^fcon3/fcon3^ (Pcdh-γ mutant,magenta) and Gad67-GFP (Pcdh-γ WT, green) cells. The total numbers of surviving cIN were counted at 6 and 21 DAT throughout the cortex. **B.** Photographs of representative coronal sections of transplanted Pcdh-γ WT (control) Ai14 labeled cells alone at 6 and 21 DAT. Note the decrease of surviving cIN between 6 and 21 DAT. Scale bar 100um. **C.** Absolute number of surviving Ai14 labeled Pcdh-γ WT cIN at 6 and 21 DAT (above). A 41.5% drop in cIN number was observed between 6 and 21 DAT (below). **D.** Photographs of representative coronal sections of transplanted Ai14 labeled Pcdh-γ mutant (magenta) and GFP labeled Pcdh-γ WT (green) cells at 6 and 21 DAT. Survival of the cIN drops for both genotypes, but the Ai14 labeled cells were nearly depleted by 21 DAT. Scale bar 100um. **E.** Absolute number of surviving cIN at 6 and 21 DAT (above). Comparing 6 and 21 DAT a drop of 62.8% and of 96.0% was observed, respectively, for cells WT and mutant for Pcdh-γ (below). ANOVA, F= 128.65[age], ****p<0.0001; F = 9.74[genotype], **p<0.0066, n=5 animals per time point.

### Loss of Pcdh-γ isoforms γC3, γC4 and γC5 is sufficient to increase cell death

Our results indicate that the loss of function of all 22 Pcdh isoforms encoded from the Pcdh-γ cluster significantly increased cell death among cINs. Whether all 22 Pcdh-γ are equally involved in the regulation of cIN survival remains unclear. Our qPCR expression analysis suggests that the expression of Pcdh-γ A1, A2, C4, and C5 in cINs increases during, or soon after, the period of cell death (**Figure 1C**). To test if Pcdh-γ A1, A2, and A3 were required for the normal survival of MGE-derived cINs, we crossed the Pcdh-γ^gAko/gAko^ mouse line (Pcdh-γ A1-A2-A3 mutant) to Nkx2-1^Cre^; Ai14 mice (**Figure 7A**). At P30, the density of cINs in both somatosensory and visual cortex of Nkx2-1^Cre^; Ai14 Pcdh-γ A1-A2-A3 mutant mice was not significantly different from that of control Nkx2-1^Cre^; Ai14 Pcdh-γ WT mice (**Figure 7D**). Consistently, co-transplanted E13.5 Pcdh-γ A1-A2-A3 mutant MGE cells and Pcdh-γ WT (GFP+) displayed surviving fractions of similar sizes (**Figure 11A, B & B’**). Therefore, the removal of the first three isoforms (A1, A2, and A3) of the Pcdh-γ cluster did not significantly affect cIN survival.

We next tested if removal of the last three isoforms of the Pcdh-γ cluster (γC3, γC4, and γC5) affected cIN survival. Pcdh-γ^gCko/+^ mice, heterozygous for the Pcdh-γ C3-C4-C5, were crossed to the *Nkx2-1^Cre^; Ai14* mouse line to label MGE/POA-derived cINs (**Figure 11A)**. While homozygous Nkx2-1^Cre^;Ai14;Pcdh-γ^gcko/gcko^ mice develop normally (normal weight and no evidence of brain abnormalities) and are born in normal Mendelian ratios, these mice die shortly after birth (Chen *et al*., 2012). To bypass neonatal lethality, and study the role of γC3, γC4, and γC5 isoforms during the normal period of cIN programmed cell death, we co-transplanted Pcdhγ C3-C4-C5 homozygous mutant E13.5 MGE cells and Pcdh-γ WT (GFP+) into the cortex of WT neonatal recipients (**Figure 11A**). At 6 DAT, the dispersion and density of tdTomato+ and GFP+ cells was indistinguishable. However, the number of tdTomato+ MGE-derived cINs dropped dramatically between 6- and 21 DAT, compared to the Pcdh-γ WT (GFP+) population (**Figure 11C**). The survival of Pcdh-γ C3-C4-C5 mutant cIN at 21 DAT was strikingly similar to that observed after transplantation of MGE cells lacking the entire Pcdh-γ-cluster (Nkx2-1-cre; ai14; Pcdh-γ^fcon3/fcon3^); compare **Figure 8 and Figure 11**. These results indicate that unlike Pcdh isoforms γA1, γA2, and γA3, Pcdh isoforms γC3, γC4, and γC5 are essential for cIN survival.

**Figure 11.**
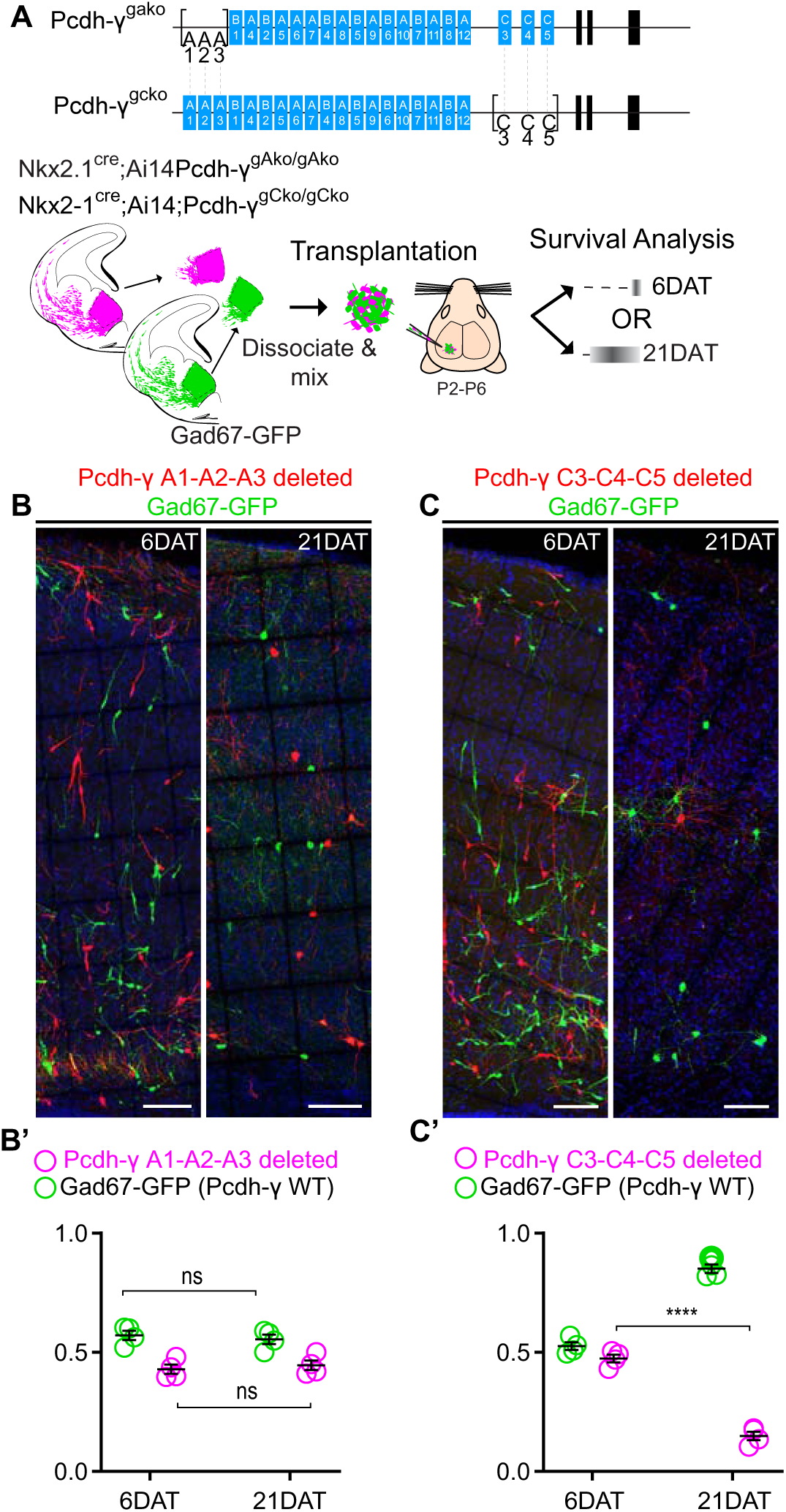
**Loss of Pcdh-γ C3, C4 and C5 is sufficient to increase cIN cell death. *A.*** (top)Diagram of mutant alleles for Pcdh-γ^gako^ (Pcdh-γ A1-A2-A3 deleted) or Pcdh-γ^gcko^ (Pcdh-γ C3-C4-C5 deleted). (bottom) Schematic of transplantation of MGE cIN precursors from Pcdh-γ A1-A2-A3 deleted (Nkx2-1^Cre^;Ai14;Pcdh-γ^gako/gako^) and Pcdh-γ C3-C4-C5 deleted (Nkx2-1^Cre^;Ai14;Pcdh-γ^gcko/gcko^) embryos. These cells were mixed in equal proportions with MGE cells from Gad67-GFP embryos (Pcdh-γ WT, green) and transplanted into host recipients. Survival of the GFP and Ai14 labeled cells was analyzed at 6 and 21 DAT. ***B, B’.*** Representative photographs of cortical sections from transplanted host animals at 6 (left) and 21 (right) DAT. Note the similar proportions of Pcdh-γ WT (GFP+) and Pcdh-γ A1-A2-A3 deleted cells (Ai14+) at 6 and 21DAT. ***B’***,Quantifications of the survival fraction of the GFP (green) and Ai14 (Magenta) labeled MGE-derived cells at 6 and 21 DAT. Note, survival fraction remains similar and constant for both genotypes (Pcdh-γ WT and Pcdh-γ A1-A2-A3 deleted cells) between 6 and 21 DAT. Scale bars, 100um. ***C, C’***. Representative photographs from coronal brain sections of transplanted host animals at 6 (left) and 21 (right) DAT. Survival of MGE-derived cIN from Pcdh-γ C3-C4-C5 deleted embryos (red) is markedly different from MGE-derived cIN from Pcdh-γ WT embryos (GFP). Scale bars, 100um. **C’**, Survival fraction at 6 and 21 DAT of the Pcdh-γ WT (GFP) and Pcdh-γ C3-C4-C5 deleted cells (Magenta). Scale bars, 100um. ****p< 0.0001

### Morphological and Physiological maturation of cINs lacking Pcdh-γ

The above results indicate that cINs lacking Pcdh-γ genes have increased cell death, specifically when the transplanted cells reach an age equivalent to that of endogenous cINs undergoing their normal period of programmed cell death. Thus, we asked whether the loss of Pcdh-γ in cINs affected their morphological maturation during this period. We first determined the survival fraction for co-transplanted control Gad67-GFP (Pcdh-γ WT) and Nkx2-1^Cre^; Ai14; Pcdh-γ^fcon3/fcon3^(Pcdh-γ mutant) MGE-derived cIN precursors at two-day intervals during the intrinsic period of cIN cell death in the transplanted population (6, 8, 10 and 12 DAT). When equal proportions of Pcdh-γ WT and Pcdh-γ mutant cells were co-transplanted, their survival fraction remains similar up to 6 DAT, but the proportion of the Pcdh-γ mutant cells drops steadily throughout the period of cell death (**Figure 12B**). Morphological reconstructions of the transplanted cells during this period of cIN programmed cell death (**Figure 12A**) revealed no obvious differences in neuritic complexity, including neurite length (**Figure 12C**), number of neurites (**Figure 12D**), number of nodes (**Figure 12E**) and number of neurite ends (**Figure 12F**), between Pcdh-γ mutant and control Pcdh-γ cells. These results suggest that Pcdh-γ genes do not play a major role in the morphological maturation of cINs during the period of cIN death.

**Figure 12.**
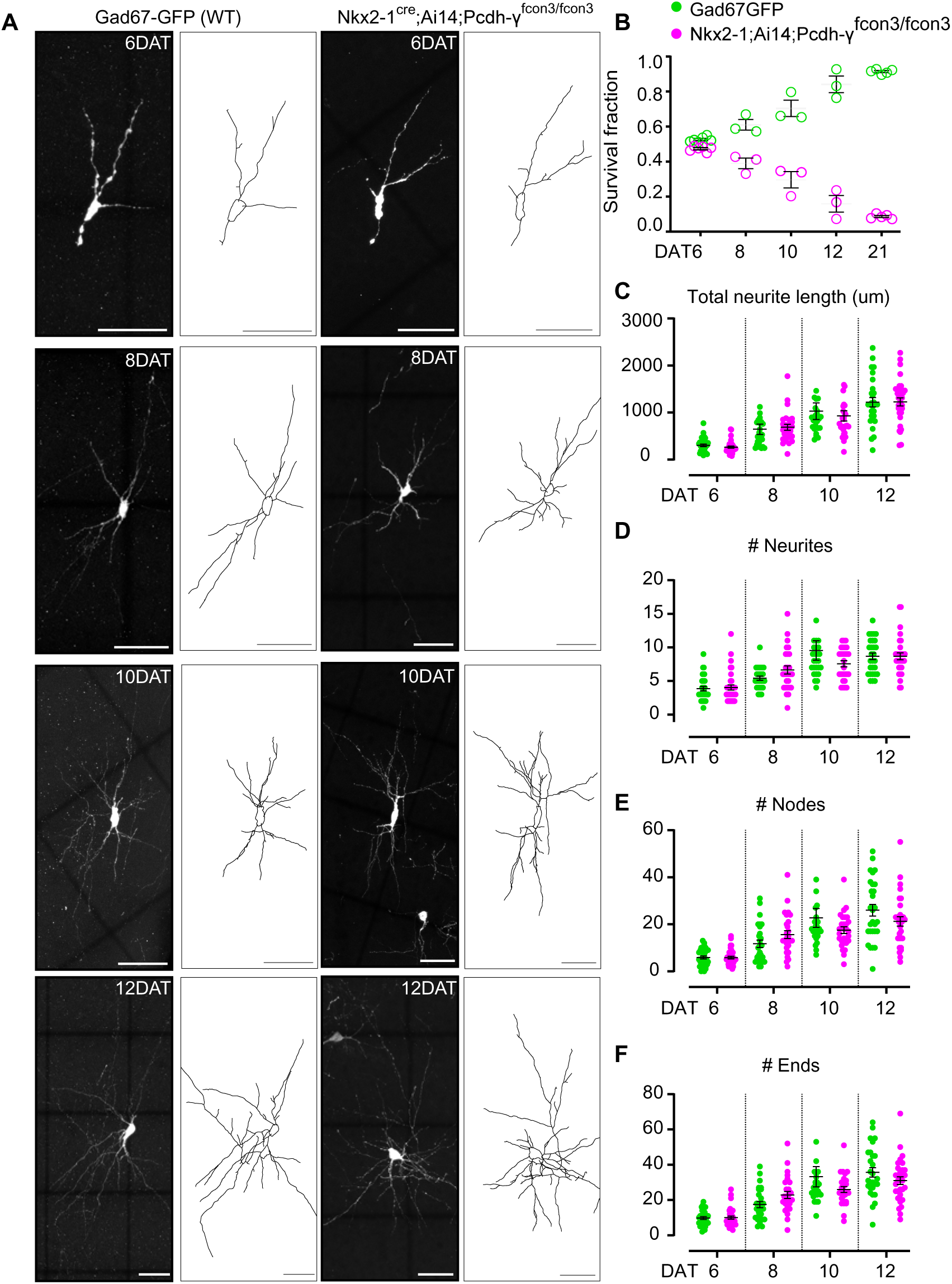
**Loss of Pcdh-γ does not affect the morphological maturation of cINs during the period of programmed cell death**. A. Photographs of representative images and morphological reconstructions of co-transplanted Gad67-GFP cells (Pcdh-γ WT, left columns) with Nkx2-1^Cre^;Ai14 Pcdh-γ^fcon3/fcon3^ cells (Pcdh-γ mutant, right columns) at 6, 8, 10 and 12DAT. Scale bars, 50um. B. Quantifications of Pcdh-γ WT(green) and Pcdh-γ mutant cells (magenta) from co-transplanted animals, represented as survival fraction from total number of cells per section at 6, 8 10 12 and 21 DAT. Pcdh-γ mutant cells begin to increase their elimination between 6 and 8DAT and this increased death occurs through 21DAT. C-F Measurements of neurite complexity during the period of programmed cell death, including neurite length (C), neurite number (D), node number (E) and neurite ends (F) Pcdh-γ WT(green) and Pcdh-γ mutant (magenta) neurons at 6, 8, 10 and 12 DAT. Two-tailed Student’s unpaired t-test, for 6DAT n= 32 cells; for 8DAT n=27 cells, for 10DAT n= 25 cells; for 12 DAT n=27 cells. All statistical comparisons were not significant following Benjamini-Hochberg multiple comparisons correction at alpha of 0.05.

Next, we utilized co-transplantation of cINs that were either Pcdh-γ deficient (Nkx2-1^Cre^; ai14; Pcdh-γ^fcon3/fcon3^) or WT (GFP+) and investigated whether the loss of Pcdh-γ affected the integration or the intrinsic neuronal properties of these cells at time points around the peak of Pcdh-γ-mediated cell death. First, to test how integration was affected, we made acute cortical slices of mouse V1 at 8, 9, 10, 11, and 12 DATs, and the frequency of spontaneous excitatory (glutamatergic) and inhibitory (GABAergic) synaptic events were recorded from the co-transplanted cINs within the same slice. There was no effect of Pcdh-γ loss on the frequency of spontaneous excitatory (glutamatergic) synaptic events or on the frequency of spontaneous inhibitory (GABAergic) synaptic events (**Figure 13B** and **Tables 1 and 2**). We next investigated whether the loss of Pcdh-γ altered intrinsic neuronal properties in co-transplanted cINs. There was no effect of the loss of Pcdh-γ on the max firing rate (**Figure 13C** and **Tables 1 and 2**), membrane time constant (Tau) (**Figure 13D** and **Table 1**), or on the input resistance (**Figure 13F** and **Table 1**). A difference in capacitance was observed between WT and Nkx2-1-cre; ai14; Pcdh-γ^fcon3/fcon3^ cINs at 8DAT, but this difference was not statistically significant following multiple comparisons correction and was not seen at later time points (**Figure 13E** and **Table 1**). We conclude that the synaptic integration, morphological and functional maturation of cINs lacking Pcdh-γ function is similar to that of WT controls.

**Figure 13.**
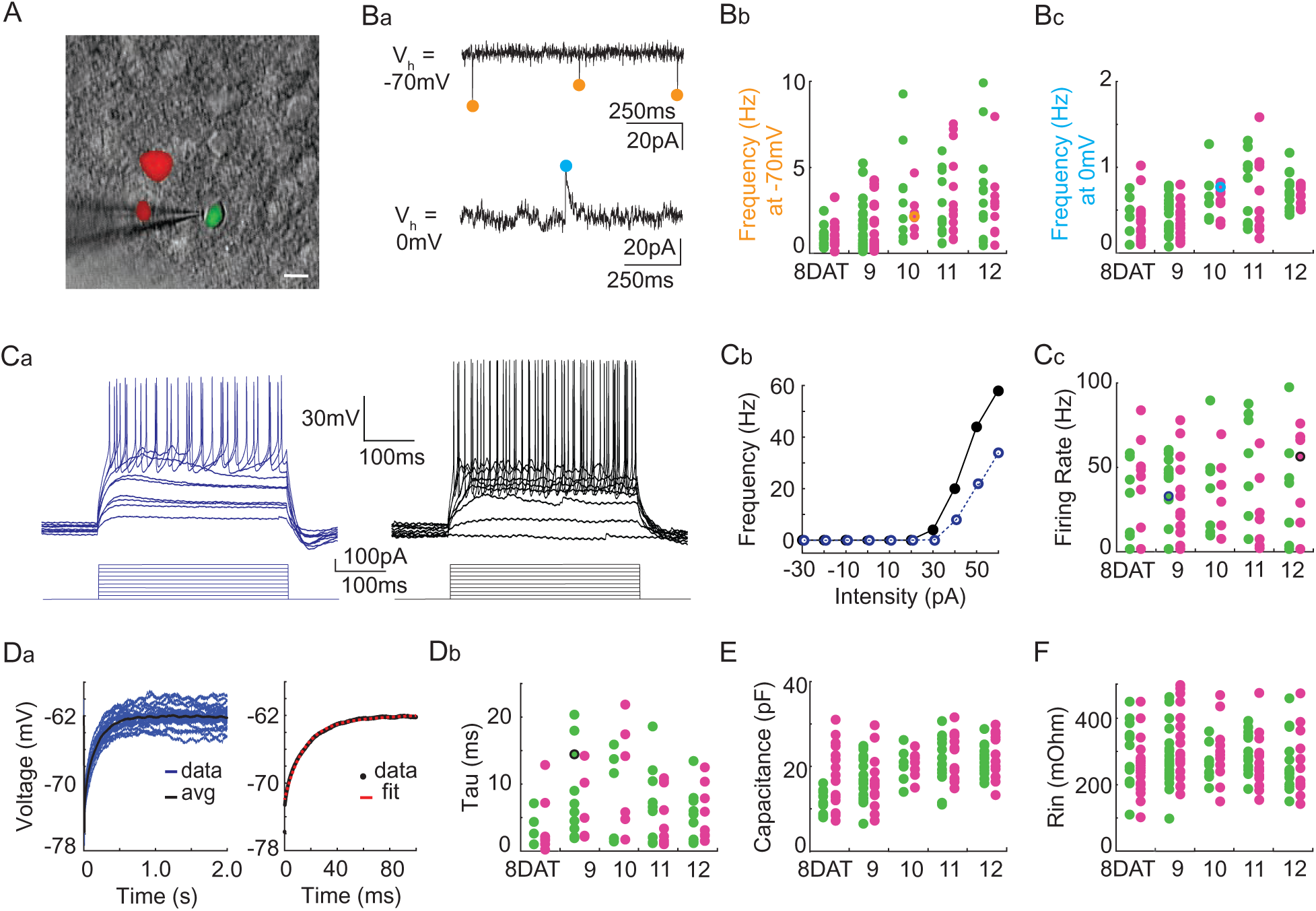
Pcdh-γ deletion does not affect the physiological properties of cINs during the period of programmed cell death. A. DIC image with fluorescence image overlaid showing co-transplanted cINs from the MGE of Gad67-GFP (Pcdh-γ WT,green) or Nkx2-1cre;Ai14;Pcdh-γ^fcon3/fcon3^ (Pcdh-γ mutant, red) embryos, recorded in an acute brain slice taken from visual cortex (scale bar 10µm). **Ba.** Representative voltage clamp recordings (1 sec) from a Nkx2-1cre;Ai14;Pcdh-γ^fcon3/fcon3^ (Pcdh-γ mutant) cIN held at −70mV (top) to record glutamatergic events (orange circles) and 0mV (bottom) to record GABAergic events (cyan circles). **Bb, Bc.** Group data from cINs recorded at 8, 9, 10, 11, and 12 DAT showing that co-transplanted Gad67-GFP cINs (WT, green circles) and Nkx2-1cre;Ai14;Pcdh-γ^fcon3/fcon3^ cINs (Pcdh-γ mutant, magenta circles) have similar rates of glutamatergic events (measured at −70mV, **Bb**) and similar rates of GABAergic events (measured at 0mV, **Bc**). The voltage clamp recordings from **Ba** are represented within the group data by the orange (−70mV) and cyan (0mV) circles in **Bb** and **Bc**, respectively. **Ca, Cb**. Representative current clamp traces showing a range of firing rates from a Gad67-GFP cIN (WT, blue trace) and a Nkx2-1cre;Ai14;Pcdh-γ^fcon3/fcon3^ (Pcdh-γ mutant, black trace) cIN responding to intracellular current injections (**Ca**), and the corresponding FI curves (**Cb**). **Cc**. Group data from cINs recorded at 8, 9, 10, 11, and 12 DAT showing that co-transplanted Gad67-GFP cINs (WT, green circles) and Nkx2-1cre;Ai14;Pcdh-γ^fcon3/fcon3^ cINs (Pcdh-γ mutant, pink circles) have similar maximum spike rates. The current clamp traces from **Ca** and **Cb** are represented within the group data by the blue and black circles. **Da**. Left: Gad67-GFP (WT) cIN voltage responses to repeated current injections (blue traces). Right: The membrane time constant (Tau) is calculated by fitting an exponential to the average voltage trace (black line). **Db.** Group data from current clamp recordings of co-transplanted Gad67-GFP cINs (WT, green circles) and Nkx2-1cre;Ai14;Pcdh-γ^fcon3/fcon3^ cINs (Pcdh-γ mutant, magenta circles) at 8, 9, 10, 11, and 12 DAT shows that Pcdh-γ deletion does not affect membrane time constant. The current clamp recording from **Da** is represented within the group data by a black circle. **E.** and **F.** Group data from current clamp recordings of co-transplanted Gad67-GFP cINs (WT, green circles) and Nkx2-1cre;Ai14;Pcdh-γ^fcon3/fcon3^ cINs (Pcdh-γ mutant, magenta circles) at 8, 9, 10, 11, and 12 DAT shows that Pcdh-γ deletion does not affect either capacitance (**E**) or input resistance (**F**).

**Table 1.** Group data for the frequency of spontaneous excitatory and inhibitory synaptic currents (Hz), max firing rate (Hz), tau (ms), capacitance (pF), and input resistance (mOhm) from figure 13. The mean, standard deviation, sample size, and statistical tests are reported. Comparisons were not statistically significant following Benjamini-Hochberg multiple comparisons correction at an alpha of 0.05.

|  | Frequency of Spontaneous Excitatory Synaptic Currents (Hz) |  |  | Frequency of Spontaneous Inhibitory Synaptic Currents (Hz) |  |  |
| --- | --- | --- | --- | --- | --- | --- |
|  | Pcdh-y<br>fcon3/fcon3 | WT | Mann-Whitney | Pcdh-y<br>fcon3/fcon3 | WT | Mann-Whitney |
|  | Mean ± SD | Mean ± SD | p value | Mean ± SD | Mean ± SD | p value |
| 8DAT | 1.0 ± 0.7 (n=18) | 1.0 ± 0.7 (n=9) | 0.89 | 0.4 ± 0.2 (n=18) | 0.4 ± 0.2 (n=6) | 0.463 |
| 9DAT | 1.4 ± 1.4 (n=19) | 1.6 ± 1.4 (n=21) | 0.432 | 0.4 ± 0.2 (n=15) | 0.4 ± 0.2 (n=18) | 0.347 |
| 10DAT | 2.2 ± 1.0 (n=9) | 3.4 ± 2.8 (n=9) | 0.652 | 0.6 ± 0.2 (n=9) | 0.6 ± 0.3 (n=7) | 0.859 |
| 11DAT | 3.2 ± 2.3 (n=14) | 2.5 ± 1.8 (n=13) | 0.298 | 0.6 ± 0.5 (n=11) | 0.7 ± 0.4 (n=11) | 0.711 |
| 12DAT | 2.6 ± 2.1 (n=11) | 3.3 ± 2.8 (n=14) | 0.597 | 0.7 ± 0.1 (n=10) | 0.7 ± 0.2 (n=13) | 0.436 |
|  | Max Firing Rate (Hz) |  |  | Tau (ms) |  |  |
|  | Pcdh-y<br>fcon3/fcon3 | WT | Mann-Whitney | Pcdh-y<br>fcon3/fcon3 | WT | Mann-Whitney |
|  | Mean ± SD | Mean ± SD | p value | Mean ± SD | Mean ± SD | p value |
| 8DAT | 21.8 ± 23 (n=16) | 21.5 ± 22.9 (n=12) | 0.789 | 3.6 ± 4.3 (n=8) | 3.8 ± 2.6 (n=4) | 0.683 |
| 9DAT | 29.1 ± 25.1 (n=17) | 35.0 ± 22.3 (n=19) | 0.416 | 6.8 ± 5.3 (n=5) | 8.7 ± 6.7 (n=10) | 0.717 |
| 10DAT | 35.7 ± 22.8 (n=6) | 37.7 ± 28.6 (n=7) | 1 | 11.0 ± 8 (n=6) | 7.8 ± 6.7 (n=6) | 0.387 |
| 11DAT | 24.5 ± 24.4 (n=11) | 36.3 ± 34.9 (n=12) | 0.534 | 5.7 ± 4.3 (n=10) | 7.3 ± 5.7 (n=9) | 0.707 |
| 12DAT | 35.0 ± 32.3 (n=10) | 29.8 ± 29.3 (n=12) | 0.757 | 5.8 ± 4.1 (n=8) | 5.5 ± 3.9 (n=9) | 0.88 |
|  | Capacitance (pF) |  |  | Input Resistance (mOhm) |  |  |
|  | Pcdh-y<br>fcon3/fcon3 | WT | Mann-Whitney | Pcdh-y<br>fcon3/fcon3 | WT | Mann-Whitney |
|  | Mean ± SD | Mean ± SD | p value | Mean ± SD | Mean ± SD | p value |
| 8DAT | 18.3 ± 7.3 (n=16) | 11.6 ± 2.7 (n=10) | 0.019 | 257.3 ± 87.1 (n=23) | 313.6 ± 86.5 (n=16) | 0.027 |
| 9DAT | 16.4 ± 6.0 (n=15) | 17.2 ± 5.1 (n=19) | 0.465 | 327 ± 106.5 (n=23) | 278.9 ± 83.4 (n=23) | 0.095 |
| 10DAT | 20.5 ± 2.8 (n=9) | 20.2 ± 3.9 (n=7) | 0.837 | 297.5 ± 94.3 (n=11) | 264.7 ± 52.9 (n=9) | 0.304 |
| 11DAT | 22.4 ± 5.1 (n=15) | 20.4 ± 6.5 (n=13) | 0.503 | 269.1 ± 79.6 (n=15) | 294.2 ± 59.9 (n=14) | 0.185 |
| 12DAT | 23.2 ± 5.8 (n=13) | 21.5 ± 3.4 (n=15) | 0.338 | 266.5 ± 96.4 (n=13) | 262.3 ± 83.6 (n=15) | 0.882 |

**Table 2.**
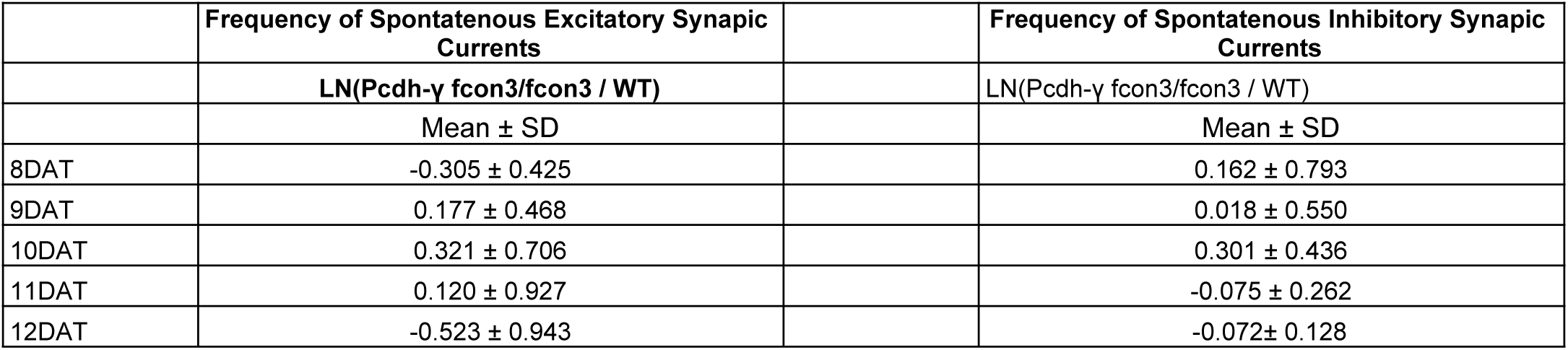
Ratios are reported for the frequency of spontaneous excitatory and inhibitory synaptic currents where wild type and Pcdh-γ ^fcon3/fcon3^ co-transplants are compared within the same slice. The natural log of each ratio was averaged for 8, 9, 10, 11, and 12 DAT and show that slice variation was not significant.

|  | Frequency of Spontaneous Excitatory Synaptic Currents |  | Frequency of Spontaneous Inhibitory Synaptic Currents |
| --- | --- | --- | --- |
|  | LN(Pcdh-γ fcon3/fcon3 / WT) |  | LN(Pcdh-γ fcon3/fcon3 / WT) |
|  | Mean ± SD |  | Mean ± SD |
| 8DAT | -0.305 ± 0.425 |  | 0.162 ± 0.793 |
| 9DAT | 0.177 ± 0.468 |  | 0.018 ± 0.550 |
| 10DAT | 0.321 ± 0.706 |  | 0.301 ± 0.436 |
| 11DAT | 0.120 ± 0.927 |  | -0.075 ± 0.262 |
| 12DAT | -0.523 ± 0.943 |  | -0.072 ± 0.128 |

## DISCUSSION

The findings above indicate that Pcdh-γ genes play a critical role in regulating cIN survival during the endogenous period of cIN programmed cell death. Specifically, the work suggests that γC3, γC4 and γC5 isoforms within the Pcdh-γ cluster are essential for the selection of those cIN that survive past the period of programmed cell death and become part of the adult cortical circuit. Pcdh-γ genes do not affect the production or migration of cINs and appear to be dispensable for the survival of cINs beyond the critical period of cell death. Together with previous work in the spinal cord and retina, these results suggest that Pcdhs γC3, γC4, and γC5 are key to the regulation of programmed cell death. In contrast, deletions of the α and β clusters did not alter cell death during this period.

Our initial approach involved the removal of Pcdh-γ function from all Gad2 expressing cells using the Gad2^Cre^; Ai14; Pcdh-γ^fcon3/fcon3^ mice. These mice displayed a dramatic reduction of cortical interneurons of all subtypes, including a significant decrease in the number of VIP+ cells, which are derived from the CGE. In these mice Pcdh-γ function was also removed from most other GABAergic neurons throughout the nervous system, as well as from a small fraction of astrocytes (Taniguchi *et al*., 2011). These general effects may, in a non-cell autonomous manner, have affected the survival of cINs in Gad2^Cre^; Ai14; Pcdh-γ^fcon3/fcon3^ mice. We therefore removed Pcdhs-γ function specifically from Nkx2-1 expressing cells in our *Nkx2-1^Cre^; Ai14;* mice. As in the Gad2^Cre^; Ai14; Pcdh-γ^fcon3/fcon3^ mice, a similar decrease in cIN was observed, but now only MGE-derived PV, SST and a subpopulation of RLN cIN were affected. The number of VIP cells was not affected in these mice, suggesting that the reduction of the number of MGE-derived cINs does not affect the survival of those derived from the CGE.

Pcdhs-γ are not required for the normal production of young cINs in the MGE, or for their migration into the mouse cerebral cortex. The extent of proliferation in the MGE was essentially the same in Pcdh-γ loss of function and WT mice, and the number of migrating MGE-derived cINs in cortex was also indistinguishable between mice lacking Pcdh-γ function and WT mice. This suggests that Pcdhs-γ are not required for the production or migration of cINs. We did not directly address whether interference with Pcdh-α and Pcdh-β isoforms affected the birth and migration of cINs, but we infer these two clusters also have no, or minimal effects on cIN production and migration, as the final number of MGE-derived cINs was not significantly affected after the loss of Pcdhs-α or Pcdhs-β. However, the aggregate loss of Pcdhs in multiple clusters may be required for phenotypes to be manifested (Ing-Esteves *et al*., 2018). We, therefore, cannot exclude the possibility that simultaneous elimination of the Pcdh-α and Pcdh-β isoforms might have an effect on cIN production, migration, or apoptosis. However, the elimination of Pcdhs-γ alone, and specifically of the γC3, γC4 and γC5 isoforms increases cell death among cINs, suggesting that removal of a limited number of isoforms in the Pcdh-γ cluster is sufficient to reveal the cell-death phenotype. Importantly, the increase in programmed cell death observed after loss of Pcdh-γ function was fully rescued when the pre-apoptotic gene Bax was also eliminated. Not only was the increased Pcdh-γ dependent cell death eliminated in Bax mutant animals, but these animals had ∼ 40% increase in the numbers of surviving cINs compared to WT controls, identical to the effect of the Bax mutation in wild type animals. This observation is consistent with previous observation showing that ∼40% of cINs are eliminated during the period of programmed cell death (Southwell *et al*., 2012; Wong *et al*., 2018). Moreover, the increased death of cINs after removal of Pcdh-γ function occurs precisely during the normal period of programmed cell death. These observations indicate that Pcdh-γ isoforms are required specifically to regulate cIN numbers during the critical window of programmed cell death. This is consistent with previous studies in retina and spinal cord that have pointed to Pcdhs-γ, and specifically the C isoforms, as key mediators of programmed cell death (Lefebvre *et al*., 2008; Chen *et al*., 2012). Interestingly, in all three neural structures, the cortex, the spinal cord, and the retina, Pcdh-γ C isoforms appear to be the key regulators of survival of local circuit interneurons. It is tempting to speculate that some or all of the C isoforms within the Pcdh-γ cluster may have evolved as important regulators of programmed cell death among local circuit interneurons. We infer that isoforms C1 and C2 in the Pcdh-α cluster are not involved in the control of cIN cell death as we did not see a cIN cell death phenotype after removal of the entire α cluster. A recent study suggests that C4 is the key mediator in the regulation of neuronal cell death in the spinal cord(Garrett *et al*., 2019). How these specific C isoforms in the Pcdh-γ cluster mediate cell death remains a fundamental question for future research. Interestingly, C4 isoform appears to be unique in that it is the only Pcdh that appears not to bind in a homophilic manner (Garrett *et al*., 2019). It is possible that this isoform has evolved as a general sensor of population size to adjust local circuit neuron numbers.

The heterochronic transplantation of cIN from the MGE into the cortex of WT mice allowed us to test for the survival of Pcdh-γ WT and Pcdh-γ mutant cIN simultaneously and in the same environment. As previously reported (Southwell *et al*., 2012), cINs die following their own time-course of maturation. Consistent with this, the transplanted cells (extracted from the MGE only a few hours after their birth) died with a delay of 6-12 days compared to the endogenous host cINs, when transplanted into P0-P6 mice. This is consistent with the notion that the cellular age of cIN determines the timing of programmed cell death (Southwell *et al*., 2012).

Interestingly, the lack of Pcdh-γ function was clearly revealed in these co-transplants, as the survival of the mutant cells was extremely low compared to that of WT cells. The dramatic reduction in the number of cIN we observed in transplanted cIN lacking Pcdhs-γ occurs precisely during the period of programmed cell death for the transplanted population. The proportion of both Pcdh-γ WT and Pcdh-γ mutant cINs decreases as a function of the distance from the transplantation site, yet the proportion of dying cells of both phenotypes remained remarkably similar at different distances from the site of transplantation. This suggests that over a wide range of densities, the programmed cell death of cIN and the effect of Pcdh-γ on this process remained relatively constant. Interestingly, the survival of WT cIN was also reduced when co-transplanted with Pcdh-γ deficient MGE-cells. This suggests that after their migration, cell-cell interactions among young cIN, is an essential step in determining their final numbers. Our work suggests that when WT cells interact with other cIN lacking Pcdh-γ function, their survival is also compromised. Homophilic Pcdh-γ, and specifically among the C isoforms, may mediate signals for survival. When the WT population is interacting with a large pool of cells of the same age that lack the Pcdh-γ, this signal may fail and these neurons may also die, therefore explaining the increase in cIN cell death observed among WT cells. The total number of interneurons, mutant + WT, may be computed among the population of young cIN before arriving in the cortex, and their final numbers subsequently adjusted through apoptosis in cortex, possibly based on this initial count. This model would be consistent with the observation that the proportion of cIN that dies scales to the initial number of grafted cells (Southwell *et al*., 2012) or to different densities according to the distance from the transplantation (**Figure 9**). These observations are consistent with a population-autonomous mechanism playing a key role in Pcdh-γ regulated cell death among cIN of the same age (Southwell *et al*., 2012).

Unlike the spinal cord where cell death is already manifested prenatally (Wang *et al*., 2002; Prasad *et al*., 2008), cIN programmed cell death occurs mostly postnatally. Since mice lacking γC3, γC4 and γC5 isoforms die soon after birth, we could not study normal cIN cell death directly in these mutant animals. We, therefore, took advantage of co-transplantation to compare the survival of cells lacking these three isoforms. The loss of cIN lacking γC3, γC4 and γC5 isoforms was identical to that when the function of the entire Pcdh-γ cluster is lost. This further suggests that these C isoforms are the key to the regulation of cell death. The co-transplantation assay, implemented in the present study, provides a powerful tool to study how the genotypes and cell-cell interactions mediate programmed cell death among interneurons. It is tempting to suggest that these cell surface adhesion proteins in cortex and spinal cord, could mediate interactions of populations of cIN arising from distant locations, with locally produced neurons to adjust final numbers. Pcdh-γ C isoforms in young cIN could also be interacting with other Pcdhs expressed by pyramidal neurons and therefore explain the adjustment in cIN number that occurs as a function of pyramidal cell number (Wong *et al*., 2018). Recent work has shown that coordinated activity of synaptically connected assemblies of cIN and pyramidal cells is essential for the survival of MGE-derived cIN (Duan *et al*., 2020). It is possible that all or some of the Pcdh-γ C isoforms are required for cIN of the same age to find each other and form these assemblies. Similarly, the initial connectivity with excitatory cells may be dependent on the proper expression of Pcdh-γ members. It will be interesting to investigate cIN survival after transplantation into mice carrying Pcdh mutations in pyramidal cells.

During the evolution of multiple mammalian species including that of humans, the cerebral cortex has greatly expanded in size and in the number of excitatory and inhibitory neurons it contains. Interestingly, the proportion of cIN to excitatory pyramidal neurons has remained relatively constant. Appropriate numbers of inhibitory cIN are considered essential in the modulation of cortical function. The embryonic origin of cIN, far from the cerebral cortex, raises basic questions about how their numbers are ultimately controlled in development and during evolution. Coordinated increase production of inhibitory interneurons in the MGE and CGE is an essential step to satisfy the demand of an expanded cortex (Hansen *et al*., 2013). In addition, MGE and CGE derived interneurons in larger brains require longer and more protracted migratory periods (Paredes *et al*., 2016). Interneurons arrive in excess of their final number. This is ultimately adjusted by a period of programmed cell death once the young cINs have arrived in the cortex. Here we have identified C-isoforms in the Pcdh-γ cluster as an essential molecular component that regulates programmed cell death among cINs.

The fact that a cell surface adhesion protein plays a key role in this regulation suggests that interactions with other cells, possibly other cINs of the same age (Southwell *et al*., 2012), or possibly excitatory pyramidal cells (Wong *et al*., 2018), could be part of the logic to adjust the final number of these essential GABAergic cells for proper brain function. An understanding of the cell-cell interactions that use C-isoform Pcdh-γs to regulate cIN cell death should give fundamental insights into how the cerebral cortex forms and evolves.

## Methods

### Animals

R26-Ai14, Gad67-GFP, Gad2-ires-cre (Gad2^cre^), *BAC-Nkx2-1-Cre* (Nkx2-1^cre^), SST-ires-cre (SST^cre^), PV-IRES-Cre-pA (PV^cre^), Bax^-/-^ and Bax^fl/fl^ and WT C57BL/6J breeders were purchased from the Jackson Laboratory. Whenever possible, the number of males and females was matched for each experimental condition. All protocols and procedures follow the University of California, San Francisco (UCSF) guidelines and were approved by the UCSF Institutional Animal Care Committee.

Pcdh-γ loss of function mice were obtained by crossing Pcdh-γ^fcon3/fcon3^ mice with Gad2^Cre^; Ai14; Pcdh-γ^fcon3/+^ mice, Nkx2-1^Cre^; Ai14; Pcdh-γ^fcon3/+^, PV^Cre^; Ai14; Pcdh-γ^fcon3/+^ or SST^Cre^; Ai14; Pcdh-γ^fcon3/+^. Pcdh-γ A1, A2, A3 knockout mice were obtained from crosses of Pcdh-γ^gako/gako^ mice to Nkx2-1^Cre^;Ai14; Pcdh-γ^gako/+^ mice.

Embryonic donor tissue was produced by crossing WT C57BL/6J to heterozygous mice expressing green fluorescent protein-expressing (GFP) driven by Gad67. Pcdh-γ^fcon3/fcon3^ Ai14 (tdTomato-expressing) tissue was obtained from embryos produced by crossing Nkx2-1^Cre^; Ai14; Pcdh-γ^fcon3/+^ mice with Pcdh-γ^fcon3/fcon3^ mice.

Pcdh-γ^gako/gako^ Ai14(tdTomato-expressing) tissue was obtained from embryos produced by crossing Nkx2-1^Cre^;Ai14; Pcdh-γ^gako/+^ mice to homozygote Pcdh-γ^gako/gako^ mice. Pcdh-γ^gcko/gcko^ Ai14(tdTomato-expressing) tissue was obtained from embryos produced by crossing Nk2-1^Cre^;Ai14; Pcdh-γ^gcko/+^ mice to heterozygote Pcdh-γ^gcko/+^ mice. Homozygote Pcdh-γ^gcko/gcko^ are neonatal lethal. GAD67-GFP and Nkx2-1^cre^; Ai14 offspring were genotyped under an epifluorescence microscope (Leica), and PCR genotyping was used to screen for Pcdh-γ^fcon3/fcon3^, Pcdh-γ^gako/gako^ and Pcdh-γ^gcko/gcko^ embryos in GFP or Ai14 positive animals. All cell transplantation experiments were performed using wild type C57Bl/6 recipient mice. All mice were housed under identical conditions.

### Cell dissection and transplantation

Unless otherwise mentioned, MGEs were dissected from E13.5 embryos as previously described (Southwell *et al*., 2012). The day when the sperm plug was observed was considered E0.5. Dissections were performed in ice-cold Leibovitz L-15 medium. MGEs were kept in L-15 medium at 4°C. MGEs were mechanically dissociated into a single cell suspension by repeated pipetting in L-15 medium containing DNAse I (180ug/ml). The dissociated cells were then concentrated by centrifugation (4 minutes, 800 × g). For all co-transplantations, the number of cells in each suspension (GFP+ or tdTomato+) was determined using a hemocytometer. Concentrated cell suspensions were loaded into beveled glass micropipettes (≈70-90 μm diameter, Wiretrol 5 μl, Drummond Scientific Company) prefilled with mineral oil and mounted on a microinjector. Recipient mice were anesthetized by hypothermia (∼4 minutes) and positioned in a clay head mold that stabilizes the skull. Micropipettes were positioned at an angle of 0 degrees from vertical in a stereotactic injection apparatus. Unless otherwise stated, injections were performed in the left hemisphere −1 mm lateral and 1.5 mm anterior from Lambda, and at a depth of 0.8mm from the surface of the skin. After the injections were completed, transplant recipients were placed on a warm surface to recover from hypothermia. The mice were then returned to their mothers until they were perfused or weaned (P21). Transplantation of Nkx2-1^Cre^; Ai14; Pcdh-γ^gako/gako^ was performed using frozen cells (**Figure 11B**). For such experiment, dissected MGEs from each embryo were collected in 500 uL L15 and kept on ice until cryopreserved. MGEs were resuspended in 10% DMSO in L15 and cryopreserved as *in toto* explants (Rodríguez-Martínez, Martínez-Losa and Alvarez-Dolado, 2017). Vials were stored at −80 C in a Nalgene™ Mr. Frosty Freezing Container that provides repeatable −1°C/minute cooling rate according to the manufacturer, and then transferred to liquid nitrogen for long term storage. Prior to transplantation, vials were removed from −80 and thawed at 37C for 5 minutes. Freezing media was removed from vial and replaced with L-15 at 37C. Dissociation was performed as above.

### Immunostaining

Mice were fixed by transcardiac perfusion with 10mL of ice-cold PBS followed by 10mL of 4% formaldehyde/PBS solution. Brains were incubated overnight (12-24hrs) for postfixation at 4°C, then rinsed with PBS and cryoprotected in 30% sucrose/PBS solution for 48 hours at 4°C. Unless otherwise stated, immunohistochemistry was performed on 50 μm floating sections in Tris Buffered Saline (TBS) solution containing 10% normal donkey serum, 0.5% Triton X-100 for all procedures on postnatal mice.

Immunohistochemistry from embryonic tissue was performed on 20 μm cryostat sections. All washing steps were done in 0.1% Triton X-100 TBS for all procedures. Sections were incubated overnight at 4°C with selected antibodies, followed by incubation at 4°C overnight in donkey secondary antibodies (Jackson ImmunoResearch Laboratories). For cell counting and *post hoc* examination of marker expression, sections were stained using chicken anti-GFP (1:2500, Aves Labs, GFP-1020, RRID:AB_10000240), mouse anti-Reelin (1:500 MBL, D223–3, RRID:AB_843523), rabbit anti-PV (1:1000, Swant PV27, RRID:AB_2631173), mouse anti-parvalbumin (anti-PV, 1:500, Sigma-Aldrich, P3088, RRID:AB_477329), rat anti-somatostatin (SST, 1:500, Santa Cruz Biotechnology, sc-7819, RRID:AB_2302603), anti-cleaved caspase 3 (1:400, Cell Signaling Technology, 9661L, RRID:AB_2341188), rabbit anti-phosphohistone-H3 (1:500; RRID:AB_310177), rabbit anti-nkx2-1(1:250, RRID:AB_793532).

### Cell counting

For cell density counts in the Visual and Somatosensory Barrel cortex, cells were directly counted using a Zeiss Axiover-200 inverted microscope (Zeiss), an AxioCam MRm camera (Zeiss), and using Stereo Investigator (MBF). tdTomato+ cells were counted in every six sections (300 µm apart) along the rostral-caudal axis. Cell densities were determined by dividing all tdTomato+ cells by the volume of the region of interest in the binocular visual cortex or the Somatosensory Barrel cortex for each animal. For PV, SST, RLN and VIP-positive cells in the visual cortex, cells were counted from confocal-acquired images. Cell densities are reported as number of PV, SST, RLN or VIP-positive cells per mm2. Cleaved caspase-3-positive cells were counted from images acquired on Zeiss Axiover-200 inverted microscope (Zeiss), an AxioCam MRm camera (Zeiss), and using Neurolucida (MBF). Cleaved caspase-3-positive cells were counted in the cortex of every six sections along the rostral-caudal axis for each animal. For cell counts from transplanted animals, Gad67-GFP positive cells and tdTomato-positive cells were counted in all layers of the entire neocortex. Cells that did not display neuronal morphology were excluded from all quantifications. The vast majority of cells transplanted from the E13.5 MGE exhibited neuronal morphologies in the recipient brain. GFP and tdTomato-positive cells were counted from tiles acquired on Zeiss Axiover-200 inverted microscope (Zeiss), a AxioCam MRm camera (Zeiss), and using Neurolucida (MBF). For quantification of the absolute numbers of transplanted cells in the neocortex of host recipients, cells from every second coronal section were counted. The raw cell counts were then multiplied by the inverse of the section sampling frequency (2) to obtain an estimate of total cell number. For all quantifications represented as fractions, GFP positive and tdTomato-positive cells were counted from coronal sections along the rostral-caudal axis in at least 10 sections per animals. For each section, the number of GFP or tdTomato-positive cells was divided by the total cell number (GFP + Ai14) in that section.

### Neuron Morphology analysis

Recipient brains were co-transplanted with Gad67-GFP (WT for Pcdh-γ) and Nkx2-1cre; Ai14; Pcdh-γ^fcon3/fcon3^ (mutant for Pcdh-γ) MGE cIN precursors. Transplanted cells were identified in sections (50um) stained for GFP and TdTomato and analyzed at 6, 8, 10 and 12 days after transplantation (DAT). Neuron morphology was reconstructed using Neurolucida software (MBF) from confocal image stacks taken at 1 μm intervals. Both GFP or TdTomato positive neurons were reconstructed from the same 50um section. Neuron morphometric analysis was done using Neurolucida Explorer.

### RT-PCR

Total RNA was prepared from dissected cortex of P30 C57Bl/6 mice using Trizol (Invitrogen) and reverse-transcribed by Quantiscript Reverse Transcriptase (Qiagen), using a mix of oligo-dT and random primers, and according to manufacturer’s protocol.

### Slice electrophysiology

Host animals were anesthetized and decapitated at 8, 9, 10, 11, or 12 days after transplant. The brain was removed into ice-cold dissection buffer containing (in mM): 234 sucrose, 2.5 KCl, 10 MgSO_4_, 1.25 NaH_2_PO_4_, 24 NaHCO_3_, 11 dextrose, 0.5 CaCl_2_, bubbled with 95% O_2_ / 5% CO_2_ to a pH of 7.4. Coronal slices of visual cortex (200 µm thick) were cut via vibratome (Precisionary Instruments) and transferred to artificial cerebrospinal fluid (ACSF) containing (in mM): 124 NaCl, 3 KCl, 2 MgSO_4,_ 1.23 NaH_2_PO_4_, 26 NaHCO_3_, 10 dextrose, 2 CaCl_2_ (bubbled with 95% O_2_ / 5% CO_2_), incubated at 33°C for 30 min, then stored at room temperature. Fluorescently identified transplant-derived MGE-lineage interneurons (Ai14; FCON3-/- or GFP; WT) were viewed using IR-DIC video microscopy. Whole-cell current-clamp recordings were made with a Multiclamp 700B (Molecular Devices) using an internal solution that contained (in mM): 140 K-gluconate, 2 MgCl_2_, 10 HEPES, 0.2 EGTA, 4 MgATP, 0.3 NaGTP, 10 phosphocreatine (pH 7.3, 290 mosm). Data were low-pass filtered at 2.6 kHz and digitized at 10 kHz by a 16 bit analog-to-digital converter (National Instruments). Data acquisition and analysis were done with custom software written in Matlab (Mathworks). Mann-Whitney (nonparametric) test, followed by multiple comparisons correction using the Benjamini-Hochberg stepdown method for control of false discovery rate (0.05 familywise), for determination of statistical significance for all comparisons of physiological properties.

### Statistical Analysis

With the exception of slice *electrophysiology data,* all results were plotted and tested for statistical significance using Prism 8. All samples were tested for normality using the Shapiro-Wilk normality test. Unpaired comparisons were analyzed using the two-tailed unpaired Student’s *t* test for normally distributed, and Mann-Whitney test for not normally distributed samples. For multiple comparisons analysis of one variable, one-way ANOVA with post hoc Turkey’s test was used to compare the mean of each column with the mean of every other column or Dunnett test to compare the mean of each column to the mean of the control group for normally distributed samples. For samples with non-Gaussian distribution, nonparametric Kruskal-Wallis test was performed followed up with post hoc Dunn’s test. Two-way ANOVA with post host Sidak’s test was used for multiple comparisons with more than one variable. Sample sizes were estimated based on similar studies in the literature.

## Acknowledgements.

We thank Joshua Sanes for kindly proving the Pcdh-γ-fcon3 mice. We also thank Ricardo Romero, Jose Rodrigues and Cristina Guinto for technical help. Thank you to John Rubenstein, Arnold Kreigstein, and Duan Xin for discussion of experimental data. This work was supported by NIH Grants (R01 NS028478, R01 EY02517) and generous gift from the John G. Bowes Research Fund to A.A.B; NIH Grant R01DC014101 to A.R.H., the Klingenstein Foundation to A.R.H., Hearing Research Inc. to A.R.H., and the Coleman Memorial Fund to A.R.H; NIH Grants (R01EY025174 and 5F32EY029935) to M.P.S. A.A.B. is the Heather and Melanie Muss Endowed Chair and Professor of Neurological Surgery at UCSF. M.P.S. is a recipient of the Research to Prevent Blindness Stein Innovator Award.

## Supplementary Figures

**Figure 1 - Figure supplement 1.**
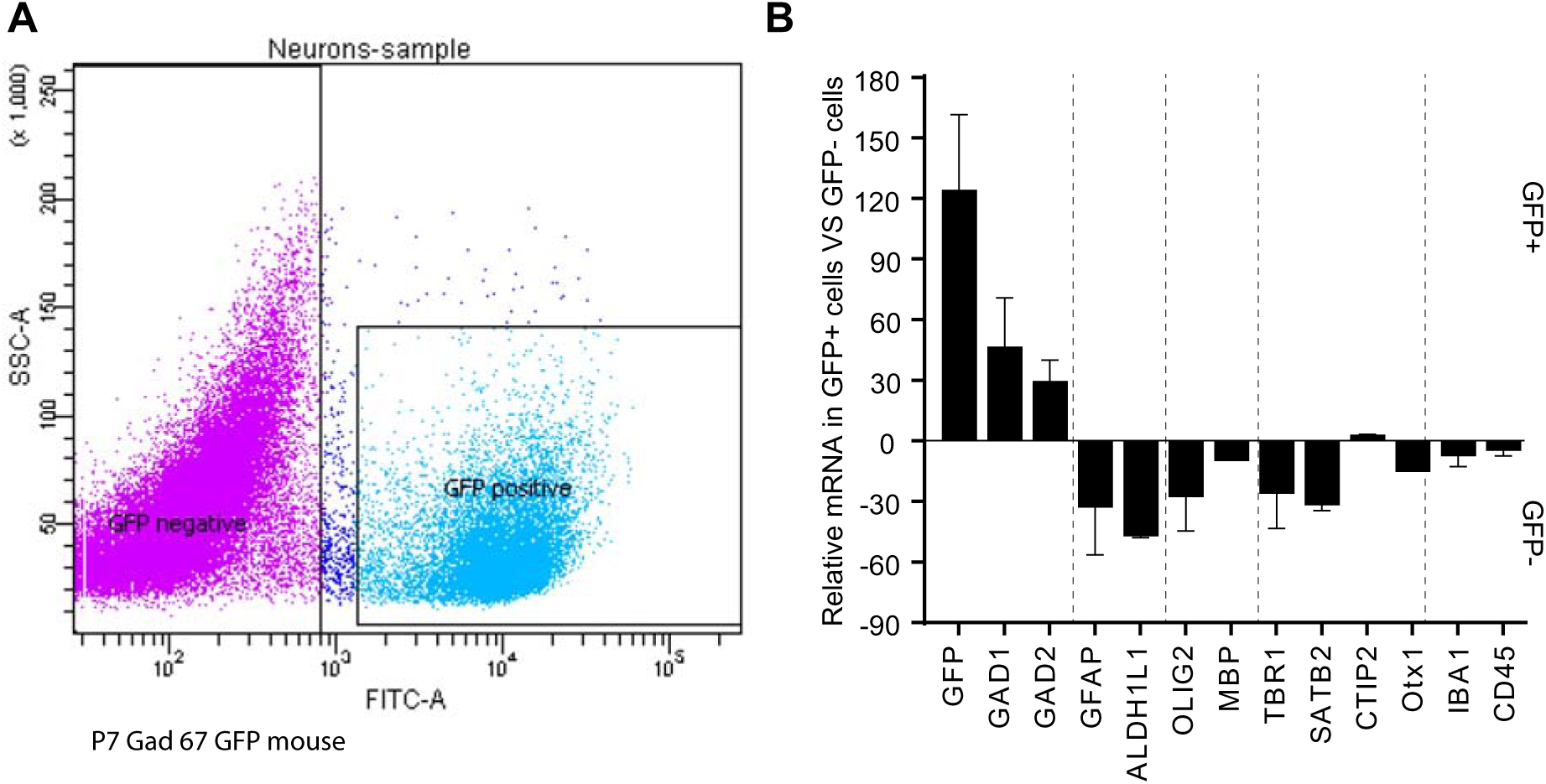
GABAergic markers are enriched in GFP positive FACS-sorted cells from Gad67-GFP mice. **A.** Representative flow cytometry plot, sorting GFP+ cIN from Gad67-GFP mice. **B.** Real-time RT-PCR analysis of GABAergic and non-GABAergic markers in positive and negative FACS-sorted GFP cells from P7 Gad67-GFP mice.

**Figure 2 - Figure supplement 1.**
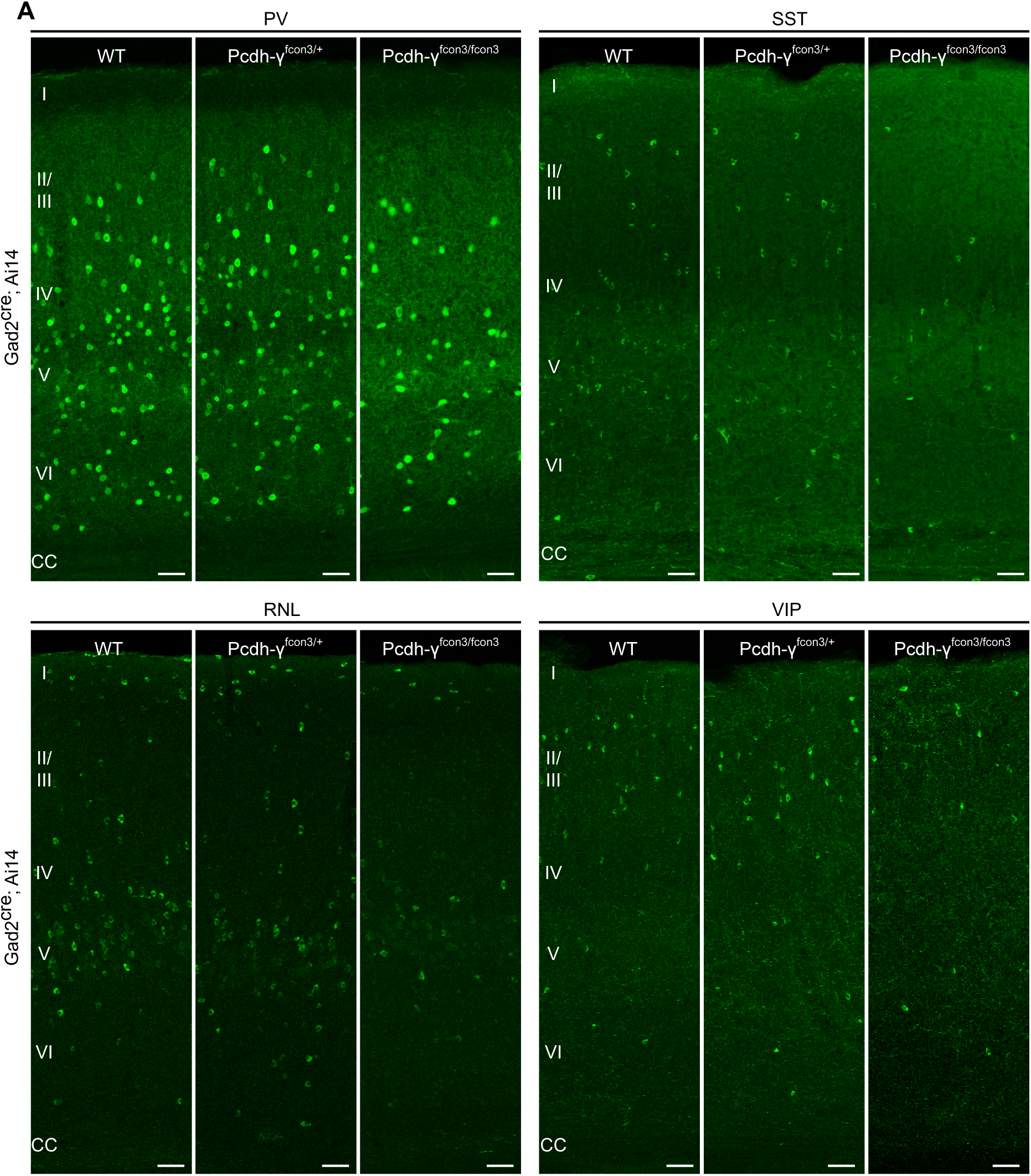
Reduced number of GABAergic cIN subtypes in γ-Pcdh deficient mice. A. Representative photographs of the binocular region of the primary visual cortex (V1b) in Gad2^Cre^;Ai14 Pcdh-γ WT, Pcdh-γ HET or Pcdh-γ mutant mice. Gad2^Cre^;Ai14 sections were stained for PV, SST, RLN or VIP. Scale bars, 50um.

**Figure 3 - Figure Supplement 1.**
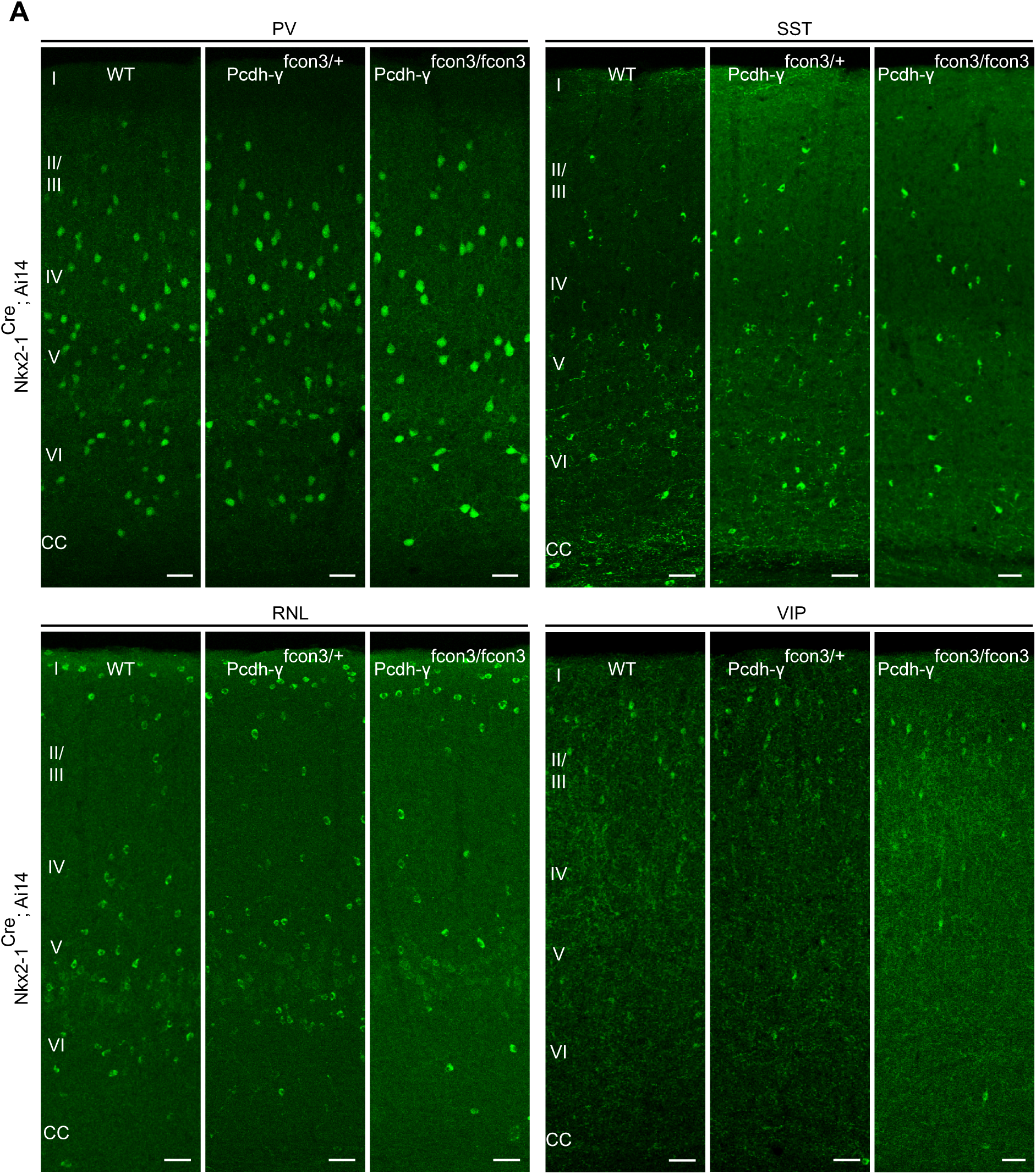
Reduced number of MGE cIN subtypes after loss of γ-Pcdh in Nkx2-1 cells. A. Representative photographs of the binocular region of the primary visual cortex (V1b) in Nkx2-1^Cre^;Ai14 Pcdh-γ WT, Pcdh-γ HET or Pcdh-γ mutant mice. Sections were stained for PV, SST, RLN or VIP. Scale bars, 50um.

**Figure 3- Figure supplement 2.**
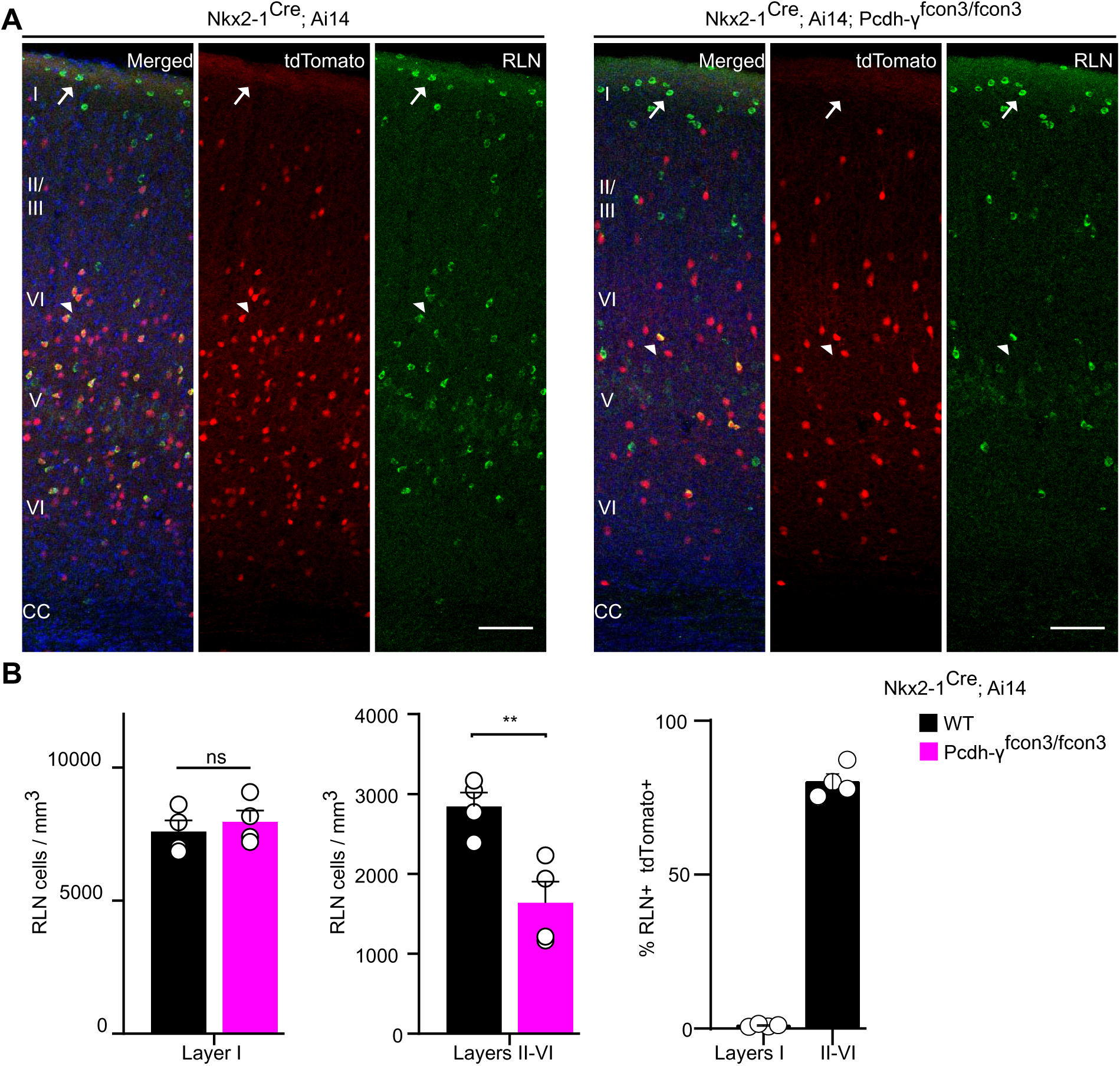
CGE-derived RLN cIN number is not affected by loss of Pcdh-γ in Nkx2-1-derived cells. A. Representative photographs of the binocular region of the primary visual cortex (V1b) in Nkx2-1^Cre^;Ai14 Pcdh-γ^+/+^v(Pcdh-γ WT, left panels) and Nkx2-1^Cre^;Ai14 Pcdh-γ^fcon3/fcon3^ mice (Pcdh-γ mutant, right panels). Sections were stained for RLN, tdTomato and DAPI. Scale bars, 100um. Arrows point to RLN positive cells, derived from CGE. Arrowhead points to RLN positive cells, derived from MGE. **B.** Quantifications of the density of RLN positive cells in layer I (left graph) and layers II-VI (middle graph) of the binocular region of the primary visual cortex. Quantifications of the percentage of Nkx2-1 derived cells by cortical layer (right graph). ANOVA, n=4 animals for each genotype; for layer I, F =0.8183, p= 0.7338; for layer II-VI, F= 9.564, **p=0.0047.

**Figure 6-Figure supplement 1.**
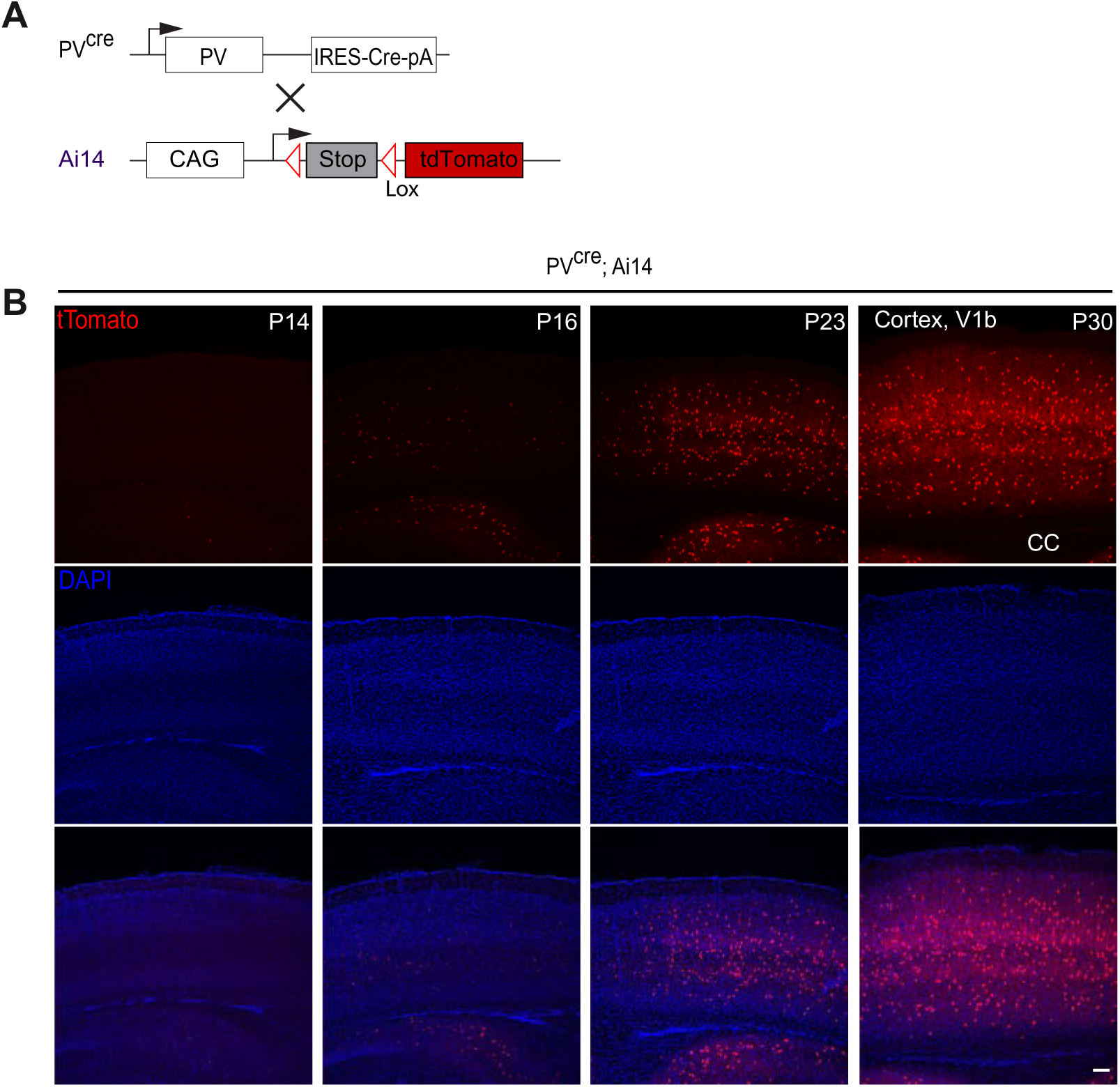
Late postnatal expression of Parvalbumin in cIN. A. PV reporter mice (PV^Cre^) were crossed to conditional Ai14 mice to label PV expressing cIN. **B.** Representative photographs of the binocular region of the primary visual cortex (V1b) in PV^Cre^; Ai14 mice at P14, 16, 23 and 30. CC= corpus callosum. Scale bars, 100um.

